# Enhancing Sampling of Water Rehydration on Ligand Binding: A Comparison of Techniques

**DOI:** 10.1101/2021.06.14.448350

**Authors:** Yunhui Ge, David C. Wych, Marley L. Samways, Michael E. Wall, Jonathan W. Essex, David L. Mobley

**Affiliations:** Department of Pharmaceutical Sciences, University of California, Irvine, CA 92697, USA; Computer, Computational, and Statistical Sciences Division, Los Alamos National Laboratory, Los Alamos, New Mexico 87545, USA; School of Chemistry, University of Southampton, Southampton, SO17 1BJ, United Kingdom; Department of Chemistry, University of California, Irvine, CA 92697, USA

## Abstract

Water often plays a key role in protein structure, molecular recognition, and mediating protein-ligand interactions. Thus, free energy calculations must adequately sample water motions, which often proves challenging in typical MD simulation timescales. Thus, the accuracy of methods relying on MD simulations ends up limited by slow water sampling. Particularly, as a ligand is removed or modified, bulk water may not have time to fill or rearrange in the binding site. In this work, we focus on several molecular dynamics (MD) simulation-based methods attempting to help rehydrate buried water sites: BLUES, using nonequilibrium candidate Monte Carlo (NCMC); *grand*, using grand canonical Monte Carlo (GCMC); and normal MD. We assess the accuracy and efficiency of these methods in rehydrating target water sites. We selected a range of systems with varying numbers of waters in the binding site, as well as those where water occupancy is coupled to the identity or binding mode of the ligand. We analyzed rehydration of buried water sites in binding pockets using both clustering of trajectories and direct analysis of electron density maps. Our results suggest both BLUES and *grand* enhance water sampling relative to normal MD and *grand* is more robust than BLUES, but also that water sampling remains a major challenge for all of the methods tested. The lessons we learned for these methods and systems are discussed.

## 2 INTRODUCTION

In their natural environment, proteins are surrounded by waters which critically affect their structure, function and dynamics.^1, 2^ Buried water molecules in the binding sites ^3–5^ also play important roles such as facilitating receptor-ligand recognition and stabilizing proteins.^2, 6–9^ A previous study done on 392 high-resolution protein-ligand crystal structures observed at least one water molecule bridging the protein and ligand in 85% of the systems.^10^

While typical MD simulations can be used to model interactions between proteins and water molecules, these often fail to adequately sample water exchange between bulk and buried hydration sites since water rearrangements in binding sites can often be extremely slow.^11, 12^ This poses significant challenges to binding free energy calculations ^13–15^ especially in relative binding free energy (RBFE) calculations which show promise in guiding experimental work in the lead optimization stage in real drug discovery projects.^13, 16^

In a typical RBFE calculation, two structurally similar ligands are compared by simulating them in both the protein-ligand complex and in the solution state where one ligand is transformed into another via unphysical or alchemical pathway. However, even closely related ligands have differences in water placement in the binding site.^17–20^ In RBFE calculations, when simulating the protein-ligand complex, the simulation timescale is normally too short (e.g., ns) to allow adequate sampling of water rearrangements when transforming one ligand to another which impairs the accuracy of such calculations.

A variety of methods seek to advance the knowledge of optimal placement of water molecules and facilitate binding free energy calculations.^21–32^ Among these methods, we are especially interested in two: nonequilibrium candidate Monte Carlo (NCMC)^33^ which efficiently hops water molecules between energy basins, and grand canonical Monte Carlo (GCMC)^34–37^ which allows the fluctuations in the number of water molecules in a simulation according to a specified chemical potential. Both methods show promise in improving water sampling in molecular simulations and GCMC has shown the ability to incorporate the thermodynamics of buried water in binding free energy calculations. ^31, 32, 38–40^ Ben-Shalom et al.^30, 41^ recently studied a Monte Carlo (MC)/MD hybrid approach and found robust sampling of buried hydration sites and improved accuracy in relative binding free energy calculations. However, this approach was implemented in a different simulation engine (AMBER package^42^) than the one used in this work (OpenMM^43^) for MD, NCMC and GCMC. So we didn’t include this approach in this work (check Section 3 for more details).

In this work, we seek to compare the efficiency and accuracy between NCMC and GCMC methods in water sampling using a broad range of systems. We also compare with plain MD simulations as a point of reference. The results from a comprehensive comparison among these techniques provide valuable lessons regarding MD simulation water-sampling issues, which are important for applications such as binding free energy calculations.

## 3 METHODS

### 3.1 Force field limitations

Before we move to simulation details, we address force field limitations, a key concern in any simulation study. Molecular simulations are conducted using an underlying energy model, or force field, which approximates the underlying physics. Even though force fields for proteins, small molecules and solvents have been developed for several decades, a perfect force field would still be an approximation, and present-day force fields still seem not to have reached the limitations of the functional form and thus are not perfect. Thus, even if all other aspects of simulations are correct (timescale, preparation, etc.) predictions from simulations still may differ from experimental measurements. In addition, other factors like the temperature at which the diffraction data was collected in experiments may also contribute to the discrepancies between simulations and experimental measurements.

In this work, when examining the efficiency of the different computational methods examined, we do not address the issue of any potential force field limitations. In general, a better sampling method ought to more efficiently yield results closer to the correct value given the chosen force field, but it won’t address force field problems (i.e., the force field does not well represent the true system or the conditions in which experimental measurements were conducted). In principle, it is possible that a better sampling method might yield worse agreement with experiment, if the correct answer for the force field differs from reality. Thus, when comparing methods, a successful simulation is one which captures the true force field answer for the system. Ideally that would also agree with experiment. But if it doesn’t, and we indeed have captured the correct force field answer for the system (which may be assessed by agreement among all of the methods examined, or with a gold standard approach, for example), in the present context we still consider such a simulation as success.

### 3.2 Selected targets

We selected the targets from two recent studies focusing on using enhanced sampling of water motions to improve the accuracy of binding free energy calculations,^30, 31^ including several proteins: Protein Tyrosine Phosphatase 1B (PTP1B), Heat Shock Protein 90 (HSP90), Bruton’s Tyrosine Kinase (BTK), transcription initiation factor TFIID subunit 2 (TAF1(2)), and thrombin. In addition to being different receptors, these targets differ in binding site positions, number and occupancy of buried water sites. We aim to include enough diversity and cover a broad range of systems which were studied previously so that we may validate our results against prior work. Several targets studied in this work also differ in the occupancy of water sites between congeneric ligands which may pose challenges in relative binding free energy calculations. Figure 1 shows the binding sites of these systems, along with crystallographic water molecules and the relevant Protein Data Bank (PDB) IDs.

**Figure 1:**
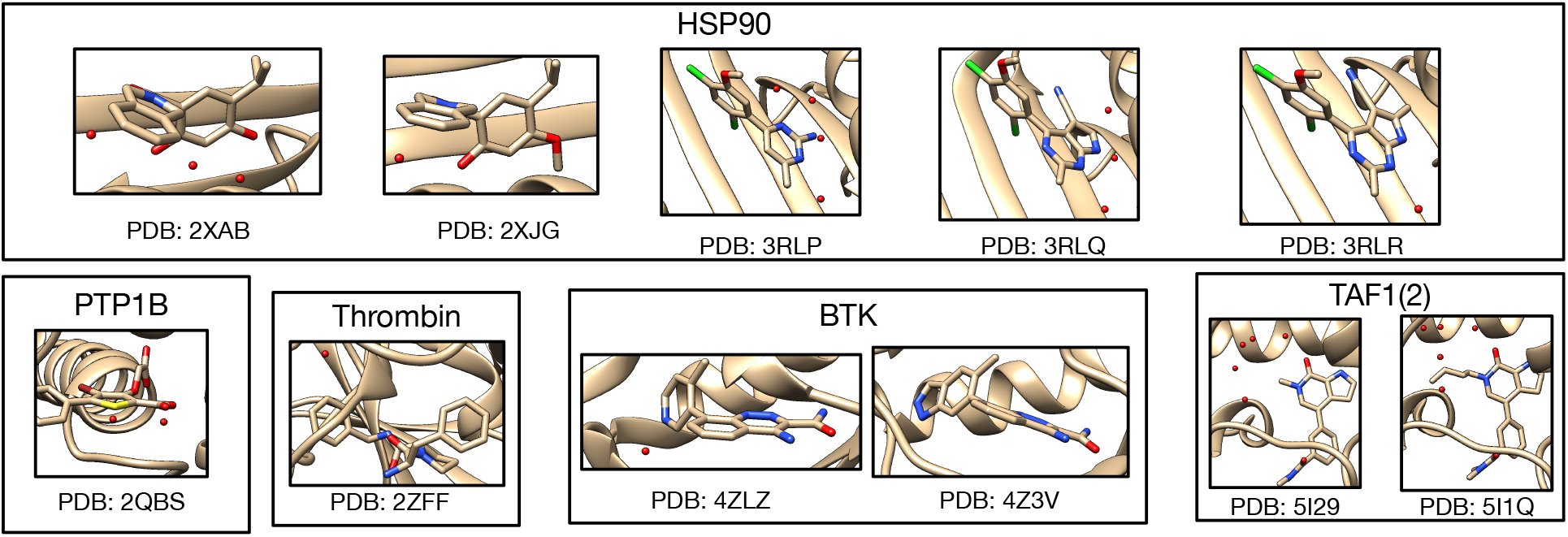
All 11 protein-ligand systems studied in this work and their hydration sites (red spheres) and their PDB IDs.

### 3.3 Molecular Dynamics Simulations

The ligand was parameterized using Open Force Field version 1.2.1 (codenamed “Parsley”).^44, 45^ The AMBER ff14SB force field ^46^ was used for protein parameterization in conjunction with TIP3P water model. ^47^ BLUES, *grand* and normal MD simulations were performed using OpenMM (version 7.4.2).^43^ A time step of 2 fs, and a friction constant of 1 ps*^−^*^1^ were used in MD simulations. Long-range electrostatics were calculated using Particle Mesh Ewald (PME)^48, 49^ with nonbonded cutoffs of 10 Å. Each system was simulated at the experimental temperature listed on the PDB website (https://www.rcsb.org). We used pdbfixer 1.6 (https://github.com/openmm/pdbfixer) to add the missing heavy atoms to the receptor. Then, the PROPKA algorithm^50, 51^ on PDB2PQR web server^52^ was used to protonate the receptors residues at experimental pH values. The pK*_a_* values of ligands were calculated using Chemicalize (ChemAxon, https://www.chemaxon.com) and then were used to determine protonation states of ligands based on the simulation pH conditions.

For each target, we performed two separate MD simulations with different starting velocities, one set (1) with ordered water molecules removed prior to simulation and another, (2) with ordered water molecules retained. Ideally, the two versions of simulations will converge to similar results (e.g., suggesting similar occupancies of target sites).

The systems first were minimized until forces were below a tolerance of 10 kJ/mol using the L-BFGS optimization algorithm ^53^ implemented in OpenMM, followed by 1 ns NVT equilibration and 10 ns NPT equilibration. The force evaluations for the two equilibration phases are 0.5 and 5 million. The production run was performed in the NPT ensemble for 70 ns of a single simulation block (equivalent to 35 million force evaluations) in consideration of our cluster’s actual wallclock time limit and was extended 9 times to 700 ns in total. Each individual 70 ns unit in this 700ns production run constitutes a simulation block for the purposes of the analysis we present here.

Our previous work showed that restraining the protein and ligand to maintain the crystallographic pose was helpful in water insertion to the target sites in BLUES simulations since this helps keep protein cavities from collapsing; when they collapse, it can be difficult for simulations to re-fill them. It is also interesting to test this idea in plain MD simulations although we believe the success of the approach might be system dependent. To do so, position restraints of 10 kcal mol*^−^*^1^ Å*^−^*^2^ were applied on the heavy atoms of the protein and all atoms of the ligand to maintain the crystallographic pose. The same simulation protocol was used as that for unbiased MD, except we applied these restraints in both minimization/equilibration and production runs. Two separate production runs were performed for 100 ns both in the presence and the absence of crystallographic water molecules.

We performed restrained MD simulations both to explore the benefits of using restraints in water sampling and to cross-validate against BLUES simulations where the same restraints were applied. As noted in Section 3.1, it is not guaranteed that the force field used in this work will produce structural results that agree with experimental structures. The restrained MD simulations can help to check if the buried hydration sites shown in the deposited crystal structures of the studied target systems are also favorable with the force field used here. We consider those hydration sites favorable in our simulations when they always were occupied in simulations or had an average occupancy of more than 70% with many transitions in simulations. This may or may not correspond to the hydration site being favorable in a thermodynamic sense, depending on whether sampling is adequate. Since both restrained MD and BLUES simulation restrained the protein and ligand to the crystallographic pose, we expect the same favorable hydration sites in both simulations. When discrepancies are observed in the results from both simulations compared to the crystal structures, it is possible that such disagreements are due to force field limitations.

### 3.4 BLUES Simulations

BLUES combines nonequilibrium candidate Monte Carlo (NCMC)^33^ with classical MD simulations to enhance the sampling of important degrees of freedom in ligand binding.^32, 54–57^ One advantage of using NCMC moves in water sampling is they can efficiently hop water molecules between energy basins and the likelihood of these moves is independent of the barrier heights which is normally a challenge in conventional MD simulations. The details of theory and implementation of BLUES in water sampling can be found in prior work.^32^

In BLUES, we defined a spherical region within which the water hops occur, using a heavy atom on the ligand which is close to the center of the ligand (selected visually) as the center of this sphere (see https://github.com/MobleyLab/water_benchmark_paper for a detailed list of selected atoms). This region should ideally cover the target water sites and extend out to bulk water to allow bulk water to exchange with water in the binding site. Since the size of the region is a parameter that may affect the success rate of NCMC moves to rehydrate the target water sites, we tested several radii for the sphere, typically using 0.8, 1.0 and 1.5 nm for most target systems. For several systems, we only used 1.0 and 1.5 nm to cover all target hydration sites.

A BLUES simulation consists of a number of BLUES iterations, where each iteration of BLUES is composed of an NCMC moves and conventional MD. In each NCMC move, interactions between the selected water molecule and its environment are gradually turned off, then the water molecule is randomly proposed to be moved to a new position in the predefined region before its interactions are turned back on. This approach allows the environment to relax in response to the proposed water translation, improving acceptance of moves and thereby accelerating water exchange and sampling. Here, we used the same number of NCMC steps (5000 steps) and MD steps (1000 steps) for all of the systems. For a single simulation block, in consideration of our cluster’s actual wallclock time limit, 3000 BLUES iterations were performed using hydrogen mass repartitioning scheme with 4 fs timesteps,^58^ resulting in 12 ns simulation time and 18 million force evaluations. In the analysis presented here, we performed 10 simulation blocks in total (120 ns, 180M force evaluations). The same restraints on the protein and ligand were applied in BLUES simulations as were used in the restrained MD simulations described earlier. Simulations were done both in the presence and absence of crystallographic water molecules.

### 3.5 *grand* Simulations

Grand canonical Monte Carlo (GCMC)^34–37^ shows particular promise for enhancing water sampling and facilitating binding free energy calculations.^31, 38–40, 59–62^ In the grand canonical ensemble the chemical potential (*µ*) of the fluctuating species (here, water molecules), the volume and the temperature is constant. The water molecules can be inserted (transferred from) or removed (transferred to) from the system to enhance water sampling — judicious choice of the chemical potential gives an equilibrium between the simulated system and bulk water.

In this work, the *grand* package ^63^ was used to perform GCMC moves with MD sampling using OpenMM simulation engine.^43^ We used OpenMM so that all of simulation techniques (MD, NCMC, GCMC) studied in this work used the same engine (OpenMM) which provides an opportunity to conduct a relatively fair comparison between these techniques, avoiding scenarios where implementation differences in different engines might bias the results. This is also one reason that the Monte Carlo (MC)/MD hybrid approach recently presented by Ben-Shalom et al.^30, 41^ was not examined in this work since the hybrid approach used there was implemented in the AMBER simulation package.^42^

In *grand*, as in BLUES, a GCMC region needs to be defined first. To do that, we selected two atoms (e.g., Cα) on the receptor so that the middle point between them is used as the center of a spherical GCMC region for enhanced water sampling (see https://github.com/MobleyLab/water_benchmark_paper for a detailed list of selected atoms). All target hydration sites are within this defined spherical region. The radius varies between systems and is dependent on the binding site size. Then the equilibration process was executed in three stages. The first GCMC/MD stage was to equilibrate the water distribution and involved an initial 10000 GCMC moves, followed by 1 ps of GCMC/MD (100 iterations, where each iteration includes 5 MD steps of 2 fs each, followed by 1000 GCMC moves). The second 500 ps NPT simulation was to equilibrate the system volume. The final GCMC/MD stage was to equilibrate the waters at the new system volume and involves 100k GCMC moves over 500 ps. The number of force evaluations for the three equilibration phases are: 0.1 million,0.25 million, and 0.35 million, respectively. The production simulation involved 2.5 ns of GCMC/MD (50 GCMC moves carried out every 1 ps of MD) for each single simulation block (1.4 million force evaluations) in consideration of our cluster’s wallclock time limit and was extended to 12.5 ns (5 blocks, 7 million force evaluations) in total. Unlike conventional MD and BLUES simulations, enhanced sampling (GCMC) of water molecules was carried out even in the equilibration phase and we found that this type of equilibration outperforms that done for MD and BLUES simulation in several systems (more details later).

There are two additional key parameters used in *grand* simulations: the excess chemical potential (*µ*^’^) of bulk water and the standard state volume of water (V*°*). Both parameters affect the acceptance probabilities of GCMC moves as depicted in previous work.^63^ For internal consistency in the GCMC/MD simulations, prior work suggested it was more appropriate to calculate the values of the excess chemical potential and standard state volume of water from simulations, rather than using the experimental values.^63^ The former is calculated as the hydration free energy of water, and the latter as the average volume per water molecule. The details of these calculations can be found in prior work^63^ and the calculated results at different temperatures used in this work can be found in Table S1.

In *grand* simulations, we only simulated the systems in the absence of crystallographic water molecules. Two separate runs were performed for each system. Based on our results, *grand* simulations were able to rehydrate all target water sites (check Section 3.7 for how we defined a success case) within five simulation blocks (12.5 ns, 7 million force evaluations) in most simulations (exceptions will be discussed later). If the results show different occupancies in water sites or the protein/ligand blocked the successful insertion of water molecules by GCMC moves, the same restraints on the protein/ligand as used in BLUES and restrained MD simulations were applied in *grand* simulations to try and help the results converge faster (e.g., 12.5 ns).

### 3.6 Trajectory Analysis

The simulated trajectories were analyzed using different approaches: (1) clustering-based analysis and (2) electron density calculations. For both approaches, MDTraj 1.9.4^64^ was used to align trajectories to the crystal structure.

#### 3.6.1 Clustering-based Analysis

We used several functions in *grand* package for clustering analysis. The water sites present within the predefined GCMC region (described in Section 3.5) were subjected to a clustering analysis, using average-linkage hierarchical clustering as implemented in SciPy, with a distance cutoff of 2.4 Å(the default parameter in *grand* package). This clustering essentially groups waters from different simulation frames which are considered to be the same site. For each cluster, the occupancy is calculated as a percentage, based on the number of frames in which that site is occupied by a water molecule relative to the total number of simulation frames. Note that, prior to clustering, a distance matrix of all water observations from the simulation was built. The distances between waters from the same simulation frame were set to an arbitrarily high value (~ 10^8^ Å) in order to discourage the merging of distinct water sites (such sites are considered distinct if they are more than 2.4 Åapart). This helps to make sure that distinct water sites which are simultaneously occupied in a single frame do not get clustered together. Otherwise, the sites might get merged and thus return occupancies greater than 100%. All of these operations were done using build-in functions in *grand* (v1.0.0/v1.0.1) package. An example script is available on https://github.com/MobleyLab/water_benchmark_paper. After we obtained these populated hydration sites in simulations, we performed a visual examination of these sites and compared them to the crystallographic waters to find the corresponding sites in the crystal structure.

For GCMC simulation data, extra steps were taken before clustering-based analysis. Particularly, as the GCMC implementation in *grand* makes use of non-interacting ‘ghost’ water molecules, which are used for insertion moves, these waters were first translated out of the simulation cell, such that they would not interfere with visualisation or structural analyses.

After we clustered water sites in simulations, we checked each site by calculating its distance to all target water sites in crystal structures. If the site representing the cluster is within 1.4 Åof the target site (as used in previous studies ^65–68^) then we consider it to be the same site. If the site representing the cluster is within 1.4 Åof more than one target site in the crystal structure, we compared the distances to these target sites and used the nearest site as the match, thereby avoiding matching sampled sites to more than one crystallographic water site

#### 3.6.2 Electron Density Calculations

Mean structure factors were computed from aligned MD trajectory snapshots. Structure factor calculations were performed using `xtraj.py`, a Python script distributed in the LUNUS open source software for processing, analysis, and modeling of diffuse scattering^69^ (https://github.com/lanl/lunus). `xtraj.py` combines methods in the Computational Crystallography Toolbox (CCTBX)^70^ and the MDTraj library for MD trajectory analysis ^64^ to compute the structure factor of each snapshot. The `xtraj.py` script is invoked at the Unix command line as `lunus.xtraj` when LUNUS is installed as a module in CCBTX. In `xtraj.py`, MDTraj I/O methods are used to read the trajectory in chunks that may be processed in parallel using MPI. A reference PDB structure is read in using the CCTBX I/O methods, and the atomic coordinates are replaced by those in a snapshot from the MD trajectory. Structure factors are computed from the modified structure using the `cctbx.xray.structure.structure_factor()` method and are accumulated within each MPI rank, along with a count of the number of frames processed. The global sums of the structure factors and frame counts are computed via MPI reduction, and the mean is computed as the aggregate sum of the structure factors divided by the frame count. Electron density maps were computed from the structure factors using CCP4 tools.^71^ By default the maps were normalized to have units of the standard deviation and a zero mean. Maps were computed using the `fft` method^72–74^ in CCP4, and were scaled in absolute units (electrons per cubic Angstroms) as needed using the amplitude of the structure factor at Miller indices (0,0,0) (F000) and volume values reported by `mmtbx.utils.f_000()`, `cctbx.xray.structure.unit_cell().volume()`, respectively, within `xtraj.py`. Example scripts to perform this analysis are available on https://github.com/MobleyLab/water_benchmark_paper.

To compare the experimental and calculated electron density maps, we visualized both maps using Coot molecular graphics ^75, 76^ (v0.9.4). We used a contour level of 3 sigma for calculated water electron density maps and 1.5 sigma for experimental protein/water maps across all systems. However, making this quantitative also requires calculating a metric describing density agreement, such as the real space correlation coefficient (RSCC). Our previous experience with RSCC suggests it may still need improvement as a metric, so measuring quantitative agreement is a research topic we do not address in the present work. Alternatively, one could quantitatively compare MD water peaks to crystallographic waters in a way described in prior work.^66^

### 3.7 Accuracy and Efficiency Comparison

Before we move on to Section 4, it is important to clarify our definition of a successful case in this work. In the simulation where all crystallographic water molecules were removed in the starting structures, we checked if all target sites could be rehydrated. To analyze our results, we must ask, “If the simulation is successful, how much will the water site be occupied in the final simulation?” The crystallographic water occupancy is not available from the experimental crystallography data (waters are universally deposited at 100% occupancy) which makes it more difficult to judge the simulation’s performance. In a recent study by Ross et al.,^31^ the average water occupancy of target water sites was checked over simulation times ranging from 30 ps to 1 ns, and simulations were considered successful when water molecules were observed to be present in the binding site some fraction of the time and improve calculated binding free energies of ligands, with no quantitative analysis of what occupancy should be considered “success”. In this work, we mainly focus on water sampling rather than predicting binding free energies. However, all studied water sites in this work have crystallographic water occupancies of 100%, as is typical for such crystallographic waters, even though a correct and converged simulation should not necessarily achieve 100% occupancy. Thus we used simulations (with ordered water retained in the initial structures) to find target occupancy for each system (more details below), with the thought that this would bias simulations towards the crystal structure and that if we saw a drop in occupancy, it would likely mean the true occupancy ought to be less than 100% with the chosen force field and model. In this way, we tried our best to avoid any bias in comparing these techniques since they all use the same force field in this work.

After checking all of our results we found our simulations fell into two categories depending on whether all simulation techniques converge to the same water site occupancies. For half of all simulated systems (PDBid: 2QBS, 2XAB, 2XJG, 3RLQ, 3RLR), the target hydration site occupancies converged to a value that was constant with longer simulations and independent of simulation technique. For example, in Figure S1, both BLUES and MD simulations with all ordered water retained prior to simulations converged to the same occupancy (100% in this case) (Figure S1A-B). In these cases, where results of long simulations agreed, we used the converged water occupancy as a reference occupancy to assess success of simulations of these systems. In the example above (Figure S1), 100% is used as our reference occupancy to check simulations where all ordered waters were removed initially to see if they can rehydrate the water site to an occupancy within 5% of the reference occupancy (100% in this example) to check for success (Figure S1C-D). We will highlight the reference occupancy we used for each system when we discuss our results below.

In this work, the reference occupancy is only used to compare the performance of these simulation techniques. It may or may not reflect the true experimental occupancy of the studied site. Such agreement would depend on the accuracy of the force field used, an aspect which would require further evaluation and is outside the scope of this work.

There are five systems (PDBid: 3RLP, 5I29, 5I1Q, 2ZFF) for which simulations results did not converge between different techniques, and thus no definition of a reference occupancy is possible. These discrepancies were caused mainly by the use of position restraints on heavy atoms in BLUES simulations and not in MD and some *grand* simulations(see METHODS). The longer timescale of MD simulations (700 ns) may also sample some protein and ligand slow motions that were not seen in BLUES and *grand* simulations, contributing to the discrepancies. In these cases where the occupancy results did not converge, we used the calculated electron density to assist in assessing the performance of simulations. In summary, if a clear converged occupancy could be obtained from simulations (Figure S1) then we used agreement of occupancy data as our success criterion. Otherwise, when we did not obtain clearly converged occupancies, we used the calculated and experimental electron density to assess the success of the simulations. We will discuss our success criteria for each system in sections below.

Here, for each technique, multiple separate simulations were performed and we checked all of them to find any simulations that achieve success for all target sites (Table 1). The simulation length of a single simulation block is not the same in different techniques (BLUES: 12ns, *grand*: 2.5ns, MD: 70ns) so the definition here is not perfect. However, in practice our results are not sensitive to this simulation time because the performance difference between these techniques is very large (more details in Table 2). There might be other good or better definitions of success than the one employed here; however, this one seems to suffice for our study, and we hope the field will settle on a more universal definition of success in future work.

**Table 1:**
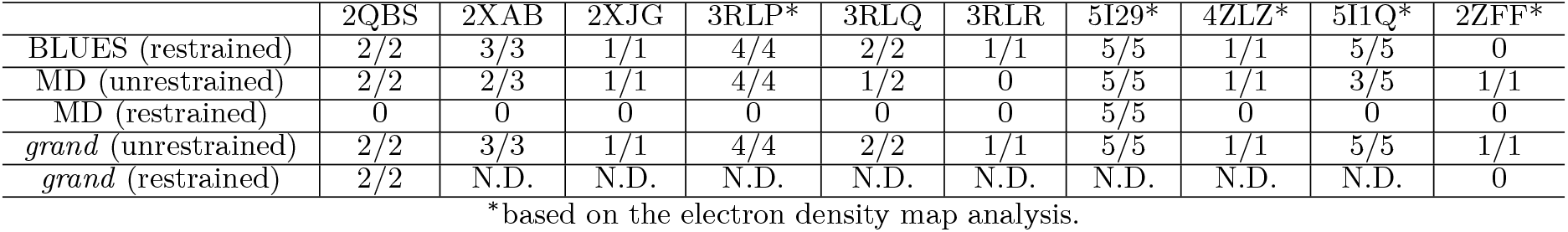
Summary of performance of each technique in each system (PDBid listed) simulation shown as n/m where n is the number of successfully rehydrated water sites and m is the number of target water sites. All ordered water molecules were removed prior to simulations. “N.D.” means no data since the simulation was not conducted. “0” indicates no water molecules were successfully inserted to the target sites.

|  | 2QBS | 2XAB | 2XJG | 3RLP* | 3RLQ | 3RLR | 5I29* | 4ZLZ* | 5I1Q* | 2ZFF* |
| --- | --- | --- | --- | --- | --- | --- | --- | --- | --- | --- |
| BLUES (restrained) | 2/2 | 3/3 | 1/1 | 4/4 | 2/2 | 1/1 | 5/5 | 1/1 | 5/5 | 0 |
| MD (unrestrained) | 2/2 | 2/3 | 1/1 | 4/4 | 1/2 | 0 | 5/5 | 1/1 | 3/5 | 1/1 |
| MD (restrained) | 0 | 0 | 0 | 0 | 0 | 0 | 5/5 | 0 | 0 | 0 |
| *grand* (unrestrained) | 2/2 | 3/3 | 1/1 | 4/4 | 2/2 | 1/1 | 5/5 | 1/1 | 5/5 | 1/1 |
| *grand* (restrained) | 2/2 | N.D. | N.D. | N.D. | N.D. | N.D. | N.D. | N.D. | N.D. | 0 |

\*based on the electron density map analysis.

**Table 2:** Summary of the efficiency of each technique (in force evaluations) in each system (PDBid listed) simulation. All ordered water molecules were removed prior to simulations. If any simulation failed to rehydrate each target site based on our defined criteria, the result is shown as “F” in the table. We calculated force evaluations (in million evaluations) required to achieve the point where all hydration sites were successfully rehydrated in each simulation. The calculated force evaluations include both equilibration and production phases. See Section 3 for detailed force evaluation in equilibration phases for each method. “N.D.” means no data since the simulation was not conducted.

|  | 2QBS | 2XAB | 2XJG | 3RLP* | 3RLQ | 3RLR | 5I29* | 4ZLZ* | 5I1Q* | 2ZFF* |
| --- | --- | --- | --- | --- | --- | --- | --- | --- | --- | --- |
| BLUES (restrained) | 8.3/13.9/6.9 | 23.5/41.5/23.5 | 6.9/6.9 | 113.5/167.5 | 41.5/23.5/23.5 | 6.9/23.5 | 6.9/6.9 | 6.9/8.3 | 12.5/13.9 | F/F |
| MD (unrestrained) | 40.5/F | F/F | 145.5/F | F/35 | F/F | F/F | 12.5/18.1 | 6.9/13.9 | F/F | 40.5/11.1 |
| MD (restrained) | F/F | F/F | F/F | F/F | F/F | F/F | 6.9/6.9 | F/F F/F | F/F | F/F |
| *grand* (unrestrained) | F/2.1 | 0.7/2.1 | 0.7/0.7 | 3.5/3.5 | 0.7/0.7 | 0.7/0.7 | 2.1/2.1 | 2.1/2.1 | 9.1/F | 2.1/2.1 |
| *grand* (restrained) | 0.7/0.7/0.7 | N.D. | N.D. | N.D. | N.D. | N.D. | N.D. | N.D. | N.D. | F/F/F |

\*based on the electron density map analysis.

When analyzing electron density maps, we must use a different criterion of success. There, we consider a test successful if the averaged electron density map calculated from simulations overlaps well with the experimental 2F*_o_*-F*_c_* map (Figure 2B) from visual inspection. In most systems, this analysis led us to the same conclusions as did the clustering based analysis. We will talk about a few exceptions later in Section 4.

**Figure 2:**
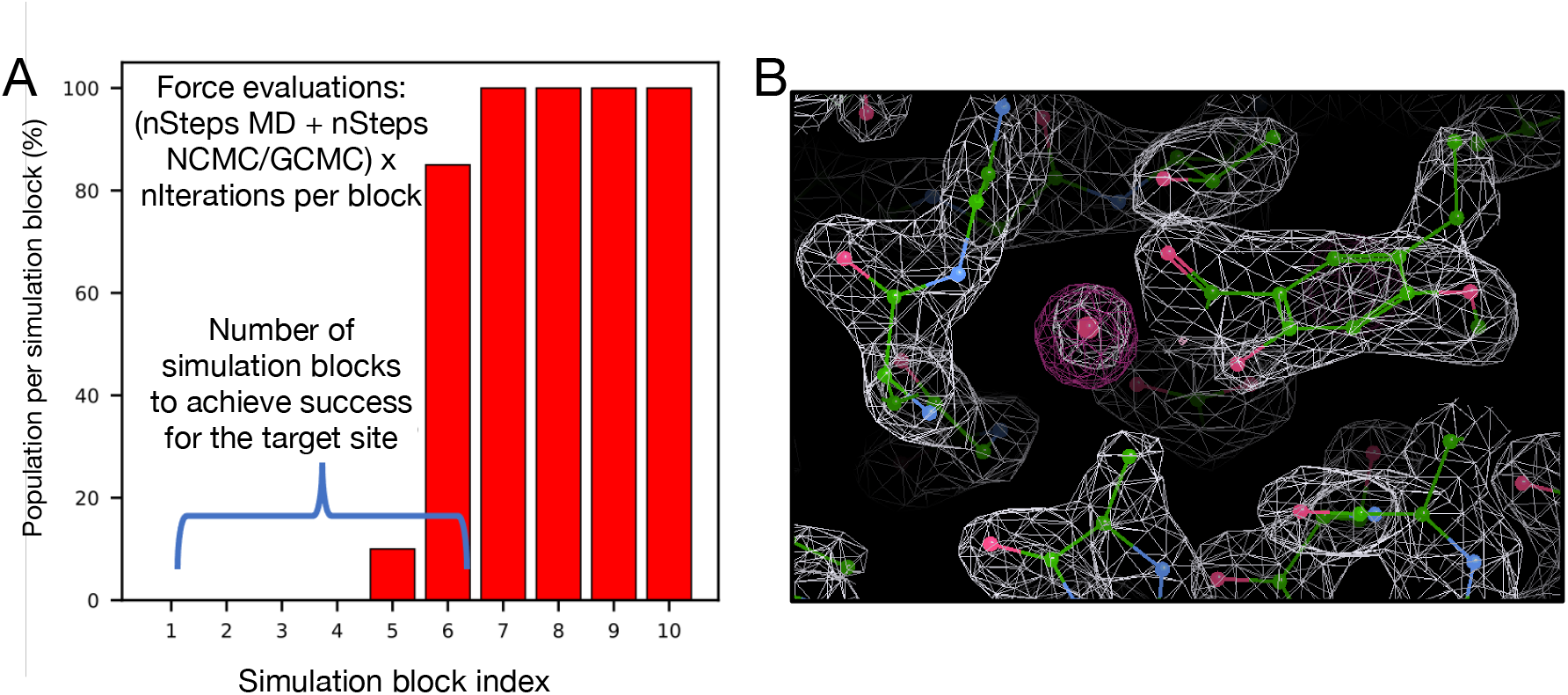
Examples of water occupancy and electron density maps for success cases. (A) Bar graphs show the water occupancy of a target hydration site in a single simulation. In this case we consider a simulation successful if the occupancy of the target water site is within 5% of 100% in a single simulation block. If successful, we check the force evaluations required to achieve this, including the number of evaluations used in equilibration. (B) The calculated electron density map of water molecules from simulation data (magenta) and the experimental determined density map (2F*_o_*-F*_c_* map) (white).

As BLUES and *grand* use both MD and NCMC or GCMC, we must account for the nontrivial cost of the NCMC/GCMC portion. Thus, we decided to use total force evaluations to compare efficiency between different simulation techniques if they successfully rehydrate all water sites based on our definitions. We did not perform efficiency analysis for failed cases (labeled as “F” in Table 2). While this is not a perfect metric, it at least does a better job accounting of these differing costs than does a more traditional metric like total simulation time. Additionally, comparisons based on wallclock time do not account for differences in compute hardware or in how much optimization has gone into improving efficiency for the particular task at hand, whereas a comparison based on force evaluations places diverse methods on relatively equal footing.

Typically, a simulation will have a total cost, in force evaluations (FEs), of (*N* + *M*) × *n* where *N* is the number of MD steps per iteration, *M* is the number of NCMC/GCMC steps per iteration, and *n* is the number of total iterations. We check the time (force evaluations) required to achieve success as defined above (Figure 2). Since multiple separate simulations were performed for each technique, we reported force evaluations required to achieve success for each simulation (Table 2). The average force evaluations across separate simulations for each technique were used when comparing the efficiency of rehydrating all target sites between these techniques. We also observed some cases where one technique was able to rehydrate all target sites in only some simulations but not all of them. In such cases, we also report the failures in Table 1 (labeled as “F”).

As noted above, the number of force evaluations in a single simulation block (in our analysis) is not the same across the different techniques (BLUES: 18 million, *grand*: 1.4 million, MD: 35 million). When the success is achieved within the first simulation block for MD and BLUES, additional analysis is performed with smaller simulation blocks so that each block has the same number of force evaluations as that of *grand* (1.4 million). In the rest of this paper, we mention the number of force evaluations for one simulation block whenever such additional analysis is performed. Except when so noted, the block length is as described in this paragraph.

GCMC moves were applied in the equilibration phases of *grand* simulations. This may introduce biases in our efficiency comparison since no enhanced sampling was applied in the equilibration phase of BLUES and MD simulations. A better way to compare these techniques in the future study would be starting simulations from the same point for production runs to exclude potential bias from starting structures equilibrated differently. However, that approach was not employed here because we chose to equilibrate with protocols which had been recommended for each method in prior work. In this work, we considered force evaluations of equilibration simulations in our efficiency comparison by reporting the sum of force evaluations of both equilibration and production phases for these methods (Table 2)

## 4 RESULTS

### 4.1 Both BLUES and *grand* outperform normal MD simulations at sampling water motions and rearrangements

Based on the definition described in Section 3.7, we calculated the overall success rate of each simulation technique.

We found that *grand* successfully rehydrated all target sites for all 10 systems (100% success rate). In contrast, BLUES failed to rehydrate the thrombin system, but succeeded in the others, giving it a success rate of 90%. The success rate was calculated by *N/M* where *N* was the number of systems that all target sites were successfully rehydrated and *M* was the number of all studied systems (*M* = 10 in this work). As we noted above, we used different success criteria for studied systems: either based on reference occupancies if possible or electron densities. But for each system, we compared different simulation techniques using the same success criterion. Both BLUES and *grand* simulations improve water sampling relative to normal MD (60% success rate) when applied to the systems studied in this work (Table 1), given the simulation lengths tested here. In those systems where all simulation techniques were able to rehydrate all target sites, normal MD proved to be much more expensive than BLUES and *grand* in most cases (Table 2).

One BTK-ligand system (PDBid: 4Z3V) studied in this work does not have buried water site and serves as a control. Particularly, we included it in this work to check if these techniques put water molecules in the binding site where no ordered water molecules were placed in the deposited crystal structures. Based on our definition described in 3 section, we do not consider this system when comparing the overall success rate between these methods. The ligand in the BTK-ligand system (PDBid: 4Z3V) observed experimentally (in crystal structures) displaces a crystallographic water molecule bridging the ligand and protein in another BTK-ligand system (PDB: 4ZLZ). In our simulations, none of the techniques employed here (MD/GCMC/NCMC) led to insertion of a water in the region from which the ligand had displaced it, confirming that all these techniques can distinguish hydration sites in the area of the binding site.

### 4.2 *grand* (GCMC/MD) is more efficient than BLUES (NCMC/MD) and MD in rehydrating all target water sites

We report the force evaluations of multiple replicates for each simulation method in Table 2. We compared efficiency using the average number of force evaluations until success across multiple replicas for each technique. In some systems, only one or two replicates of a technique were able to rehydrate all target sites. In such cases, we reported the number of force evaluations for those successful simulations and failed simulations were reported in Table 2 as “F”. Those systems where some simulations failed to rehydrate all target sites are indeed challenging for these techniques, as reflected by the fact that even in the successful simulations, it was expensive to do so (Table 2).

In all these systems, *grand* simulations more efficiently populate the water sites than BLUES or MD (Table 2). MD simulations are the most expensive in all systems. For example, in the case of a HSP90 system (PDB: 2XJG), the only successful MD simulation took 145.5 million force evaluations to rehydrate the target site whereas BLUES (6.9 million) and *grand* (0.7 million) simulations were much more efficient.

We noticed that in *grand* simulations of several targets (HSP90 (PDB: 2XAB, 2XJG, 3RLR), TAF1(2) (PDB: 5I29)), all of the targets’ water sites were rehydrated during equilibration. This is due to the fact that GCMC was used in the equilibration phase, unlike for BLUES and MD, where the equilibration was done in the normal NVT/NPT ensemble. These results highlight the benefits of using GCMC to equilibrate water molecules even without using GCMC in production runs; this approach has been shown to help obtain adequate water sampling for better binding free energy estimations even without applying it in production runs in a previous study.^31^

One potential way to take advantage of GCMC sampling in BLUES/MD simulations is running GCMC to equilibrate water molecules in prior to production runs. In this work, we did not apply this strategy in our tests. However, if all water sites are rehydrated by GCMC moves in the equilibration simulations then the approach becomes equivalent to running BLUES/MD simulations begun with all crystallographic water molecules retained, an approach we also tested in this work.

In the PTP1B system (PDB: 2QBS), it is known that maintaining the crystallographic pose is critical for successful water insertion (as observed in a previous study^31^ and elaborated via personal communication with author Greg Ross). Thus, we used position restraints on the protein/ligand (see Section 3) in *grand* simulations for this system and observed better performance compared to the unrestrained *grand* simulations (Table 2). We are able to apply restraints in this work because the crystal structures are available for all of these systems we studied. However, it is important to highlight such limitations (see additional discussion below), as these restrictions may impact *grand*’s utility in making predictions when structural information for the simulated systems might not be available.

Restraints can be used to keep a protein/ligand in a specific conformation and may accelerate the sampling of target water sites in some systems when the structure of the system along with relevant occupied water sites is known (such as from crystal structures or other techniques), as is the case here. This is an important factor we considered when we selected these systems in this study since then we can investigate the performance of these techniques in placing water in known structures. But the benefits of using such restraints are system dependent. Here, using restraints on the heavy atoms of both receptor and ligand significantly improve the performance (both efficiency and accuracy) of GCMC simulations of this PTP1B system (all three replicates rehydrated both target water sites within 2.5 ns and 1.4 million force evaluations). However, in the thrombin system (PDB: 2ZFF), using such restraints actually impairs the performance of both *grand* and BLUES (more details below). Our results also showed that only one system showed improved rehydration performance using restrained versus unrestrained MD simulations (Table 2). In contrast, a previous study found that using harmonic restraints was beneficial in the context of using MD simulations to recover the average crystallographic water structure.^66^ Such a difference is not unexpected, as the waters being studied here were removed after solvation and prior to running the simulations, whereas the previous simulations did not remove waters after solvation. Other details of the MD simulation set-up in this work and the previous study are also different (e.g., force fields, force constant for restraints, etc).

### 4.3 Lessons we learned from failures

We found that none of these simulation techniques can rehydrate all target sites in all of the systems we studied. To understand the advantages and limitations and better develop these techniques in the future, we analyze the failures.

#### 4.3.1 Failures of MD simulations

Large energy barriers can impede water rearrangements, making it difficult for unbiased MD simulation to adequately sample rearrangements of buried water molecules. Given this, we were not surprised that MD failed to rehydrate each individual target site in this study. However, we noticed that using restraints on the receptor and ligand was helpful to achieve better performance in BLUES and *grand* simulations for several targets. Thus, we tested the same restraints in MD simulations to explore potential benefits in water sampling. The results showed that no significant performance differences were observed using restraints in MD simulations compared to normal MD. We only observed in one system (TAF1(2), PDB: 5I29) in which all target sites were rehydrated faster in the simulations where restraints were applied than in unrestrained MD. In conclusion, the benefits of using restraints on the protein and ligand in using MD simulations to rehydrate vacated ordered water sites are system-dependent and are negligible in most of the systems studied in this work.

#### 4.3.2 Failures and challenges in BLUES and *grand* simulations

##### The PTP1B system (PDB: 2QBS) is a challenging case for *grand*

A previous study^31^ where GCMC was used successfully rehydrated both hydration sites in this system and we expected the same success in this work using *grand*. However, when no restraints on the protein/ligand were applied, we found that it is challenging for *grand* simulations to rehydrate Site 2 (Figure S2B). In the presence of restraints, as in BLUES, both sites could be rehydrated (Figure 3E). We used position restraints on the protein/ligand in BLUES simulations and both sites could be rapidly rehydrated (Figure 3B). These results suggest the crystallographic pose of both the protein and ligand are critical for successful insertion of water molecules in both sites, as also confirmed by the author of the previous study (personal communication) which used position restraints on the heavy atoms of the protein/ligand. ^31^

**Figure 3:**
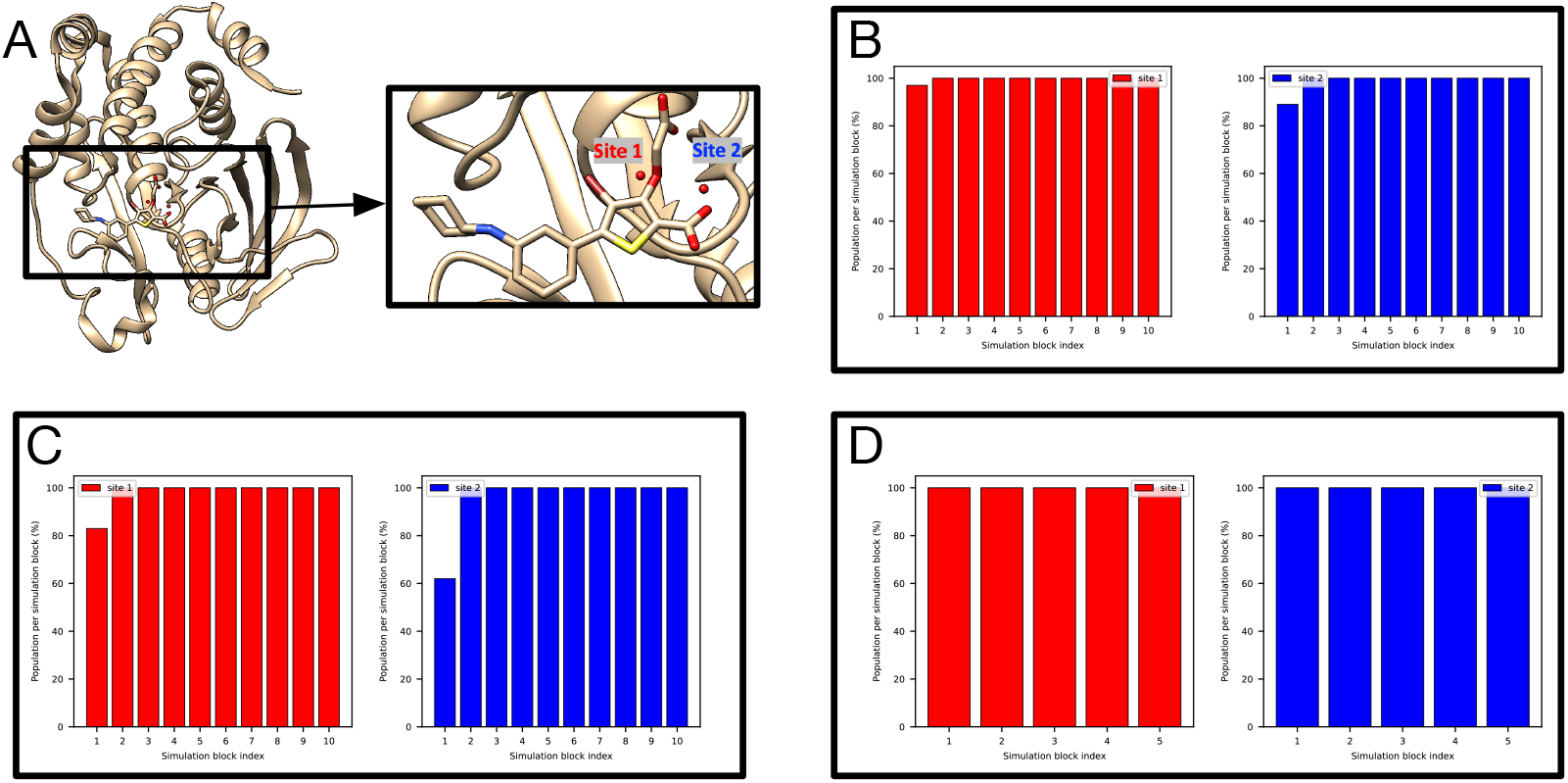
BLUES/MD/*grand* simulations can rehydrate both target sites (Site1: red, Site 2: blue) in (A) the PTP1B system (PDB: 2QBS). Bar graphs show the water occupancy of target sites in a single (B) BLUES simulation, (C) normal MD simulation, and (D) *grand* simulation. In all simulations, the ordered water molecules were removed prior to simulations. The number of force evaluations for one simulation block in (B-D) is 1.4 million, 35 million, and 1.4 million. Position restraints on the protein and ligand heavy atoms were applied in (B) and (D).

It is also worth noting that even with position restraints as used in BLUES simulations, our 100-ns simulation data show that MD simulations could not rehydrate any of the two sites. This again shows the power of enhanced sampling techniques like BLUES and *grand*.

##### The TAF1(2) system (PDB: 5I1Q) poses challenges to *grand*

This system poses challenges to *grand* simulations when the protein and ligand are not restrained; without restraints, *grand* has difficulty rehydrating Site 2 (Figure 4A). The ligand moves in the binding site when it is not restrained and may occupy the space of Site 2, blocking the successful insertion of water molecules (Figure 4A). It is also observed in MD simulations (Figure 4B). Such ligand motion is not observed in BLUES simulations when the ligand is restrained.

**Figure 4:**
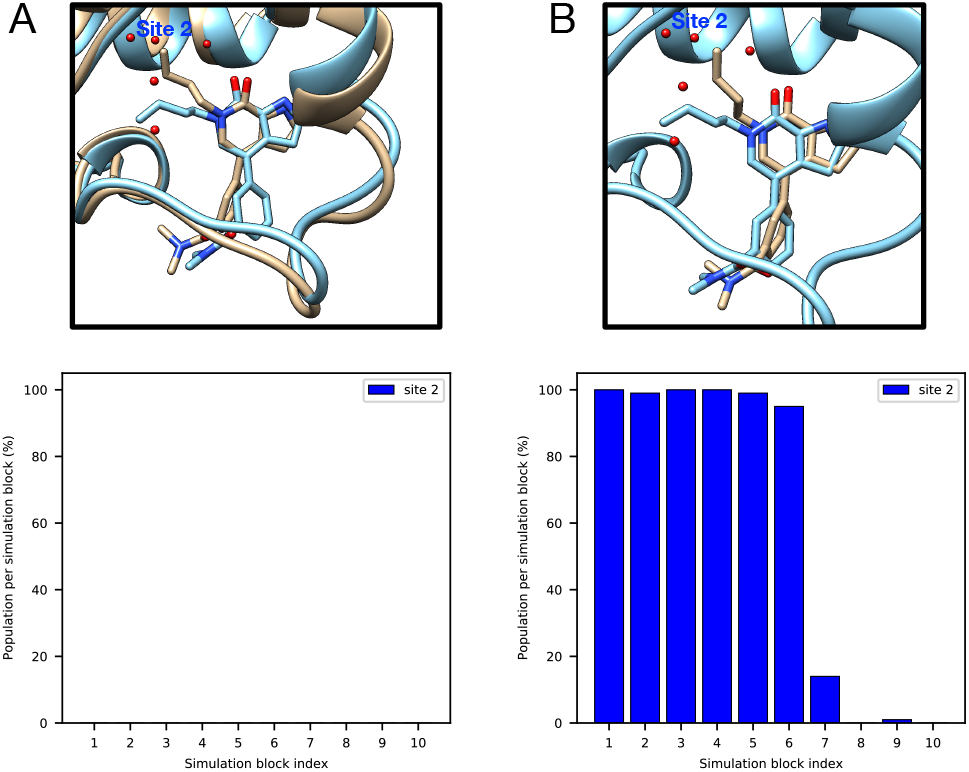
Ligand motion blocks insertion of water molecules in the target site of a TAF1(2) system (PDB: 5I1Q). Snapshots extracted from (A) *grand* simulation and (B) MD simulation. No restraints were used in either simulation. The crystallographic pose is shown in blue and simulation snapshots are shown in tan. Bar graphs show the water occupancy of Site 2 in a single (A)*grand* simulation, (B) MD simulation.

##### BLUES failed to rehydrate the target site in the Thrombin system (PDB: 2ZFF)

In the thrombin system (PDB: 2ZFF), both unbiased MD and *grand* simulations captured the target water site (Figure 5B-D). BLUES simulations, however, did not work in this system.

**Figure 5:**
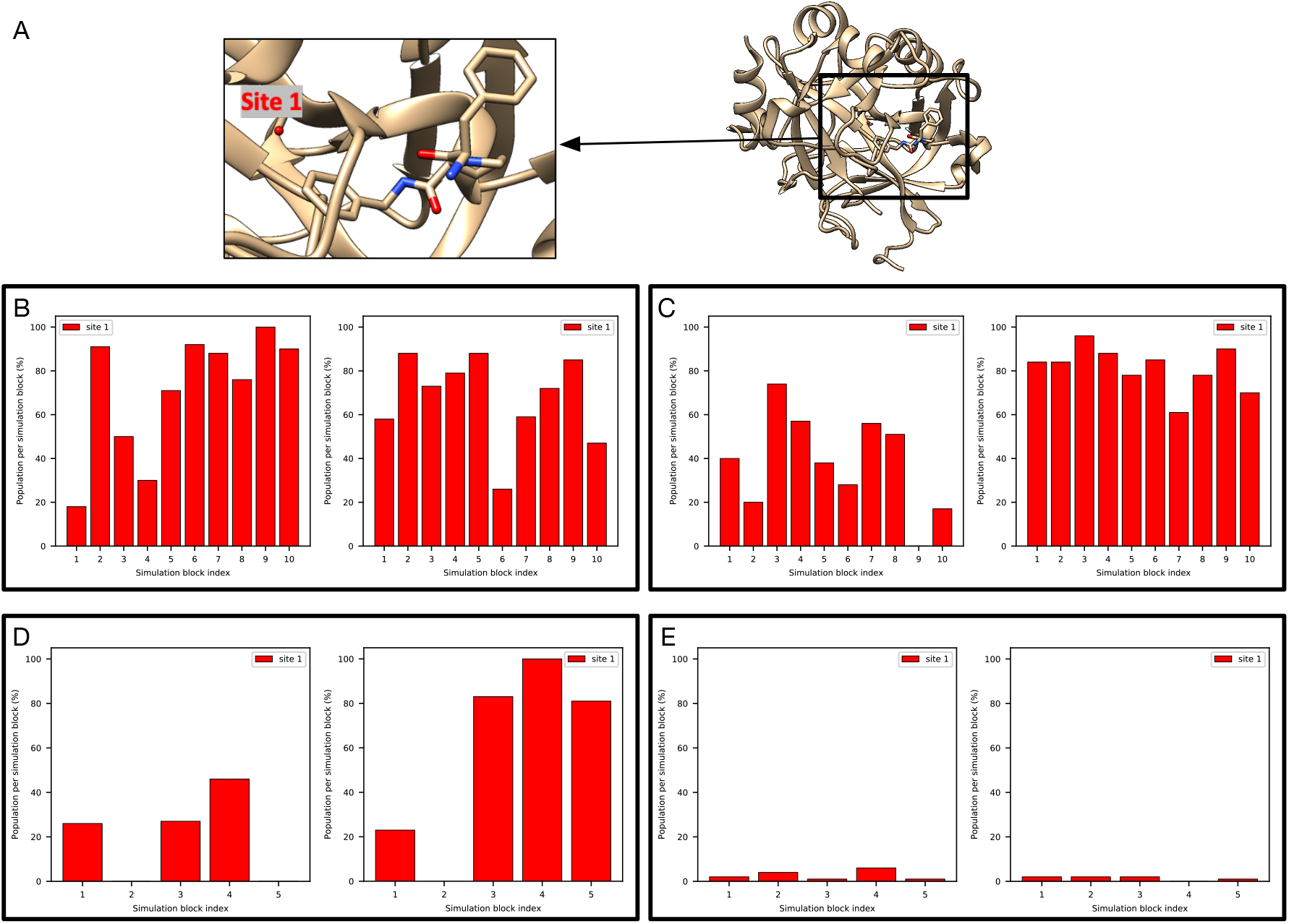
It is challenging to rehydrate the target site (red) in (A) The thrombin system (PDB: 2ZFF). Bar graphs show the water occupancy of the target site in (B) unbiased MD simulations with ordered water molecules removed prior to simulations, (C) unbiased MD simulations with ordered water molecules retained prior to simulations, (D) *grand* simulations and (E) *grand* simulations with position restraints on heavy atoms of the protein and all atoms of the ligand. All ordered water molecules were removed prior to *grand* simulations in (D-E).

The main difference between BLUES and MD/*grand* simulation protocol other than the technique itself is that restraints on the protein and ligand heavy atoms were used in BLUES simulation. We then checked the distance between selected atoms between the protein and ligand (Figure 6B) and observe a correlation between the distance and the success of water insertion (Figure 6C-F). When the distance increases, it is more likely that the water can be inserted (Figure 6C,E). In contrast, when the distance drops, the likelihood of water insertion declines (Figure 6D,F). These results suggest that additional space in the binding site is required to successfully insert the water. Thus, we find that the protein-ligand restraints used in BLUES simulations impair the performance of BLUES. We further tested this idea by using the same restraints in *grand* simulations and the probability of water insertion significantly dropped (Figure 5E).

**Figure 6:**
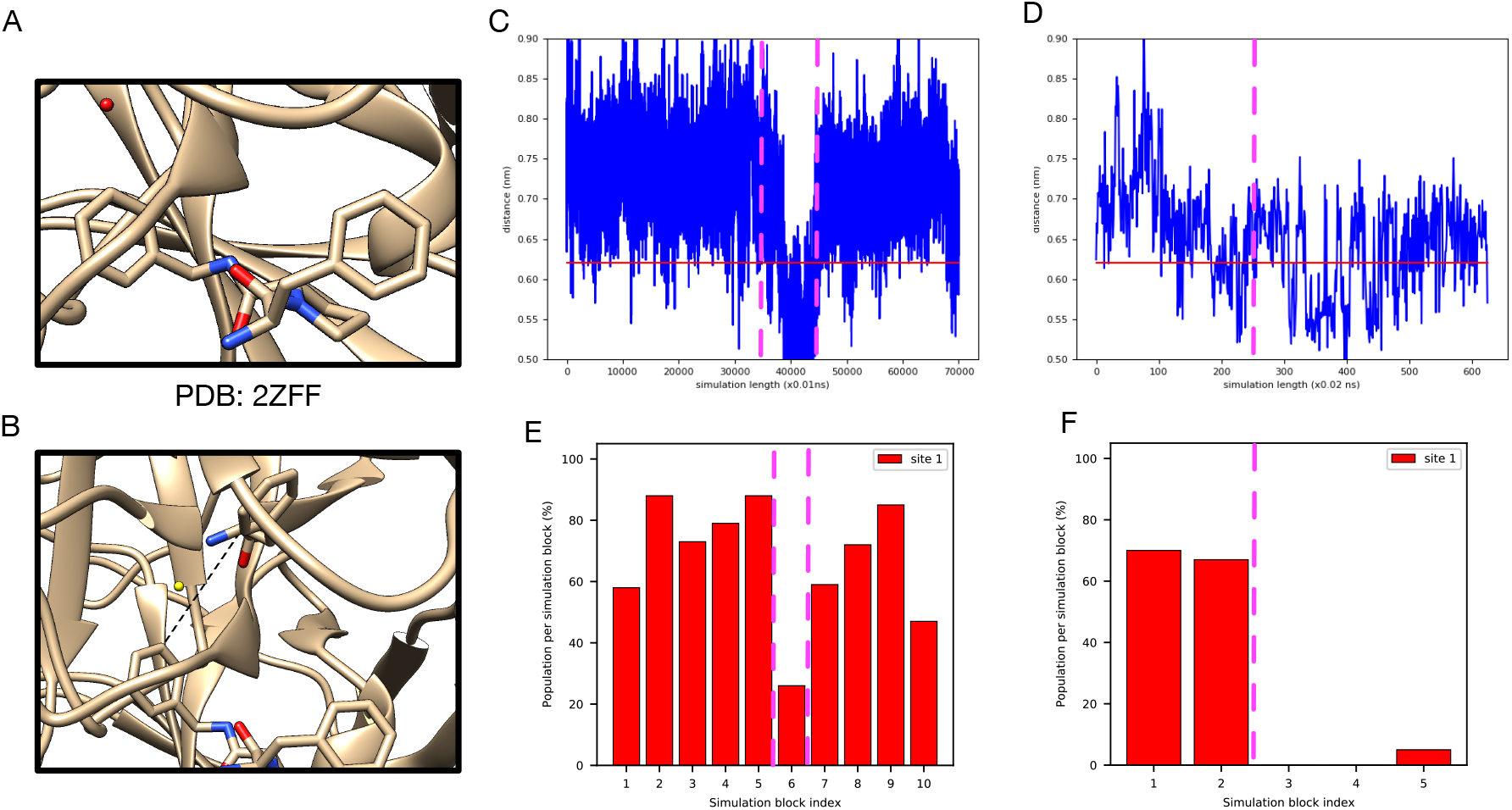
A correlation between the distance between the protein and ligand and the success of water insertion was observed in (A) the thrombin system (the target water site shown in red). (B) The atoms selected to compute distance between the protein and ligand. The distance change during (C) unbiased MD and (D) *grand* simulations (no restraints used). Bar graphs show the water occupancy of target hydration site in a single (E) unbiased MD and (F) *grand* simulation (no restraints used). The horizontal lines in (C) and (D) highlight the distance in the crystal structure (PDB: 2ZFF). The dashed vertical lines highlight the simulation block(s) in (C)-(D) and their corresponding occupancies in (E)-(F).

Since the success rate for inserting water molecules into the target site is affected by the available space between the protein and ligand, we see fluctuations in unbiased MD and *grand* simulations. It is not clear in terms of which reference occupancy to use in the case of thrombin. The reference occupancy is necessary if we want to check success using occupancies from clustering-based analysis as we discussed above. So we decide to calculate electron density from simulations and compare it with the experimental density for success check in this case.

The results are shown in Figure S3 and S4. Since BLUES simulations failed to insert water molecules in the target site, we only calculated electron density for the *grand* and MD simulations. We can see both replicates of the *grand* simulations reproduce the experimental density within the first simulation block (Figure S3). One replicate of MD simulations can reproduce the experimental density within the first simulation block but it takes much longer in the other replicate (Figure S4). Since each simulation block has different number of force evaluations for each simulation technique (MD: 35 million, *grand*: 1.4 million) in Figure and S3, we performed an additional analysis in which each simulation block has the same number of force evaluations (1.4 million) to compare the efficiency between *grand* simulations and replicate 2 of MD simulations. In Figure 7, we show the simulation block where we first observe a good agreement between the calculated and experimental electron density. We can see replicate 2 of MD simulations does not rehydrate the target site until simulation block 4 whereas both replicates of *grand* simulations can achieve it within simulation block 1. Thus, both MD replicates rehydrate the target site, but at a higher computational cost than *grand* simulations (Table 2).

**Figure 7:**
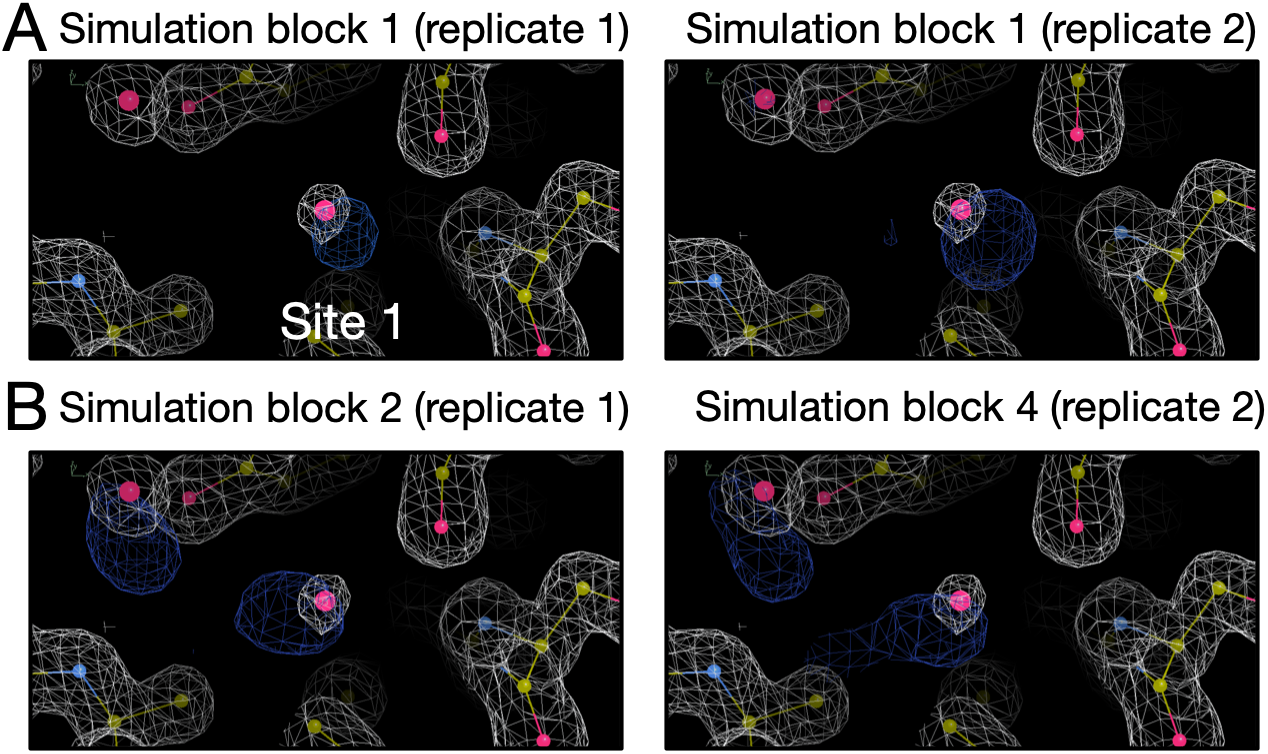
Calculated electron densities (blue) from simulations compared to the experimental densities (white) of the thrombin system (PDB: 2ZFF). The target site is labeled. We only show the simulation block when we first observe a good agreement between the calculated and experimental density. Both *grand* and MD simulations can reproduce the experimental electron densities for the target site. Panel A and B are results from *grand* and MD simulations, respectively. The left and right columns are results from two replicates for each simulation technique. The number of force evaluations for one simulation block in replicate 1 in panel B is 35 million. The number of force evaluations for one simulation block in other panels is 1.4 million.

##### All of our simulations suggest a different water network in the HSP90 system (PDB: 3RLQ) than the crystal structure

There are three target sites in this HSP90 system (Figure 8A) based on the crystal structure. However, none of our simulations could rehydrate Site 1, though they could rehydrate Sites 2-3 (Figure 8B-D). Initially, we considered this to be a failure. However, we checked simulations where all ordered water molecules were retained (as in the crystal structure) prior to simulations. We found the water molecule in Site 1 escaped quickly in simulations and Site 1 was not occupied during most of the simulations. This suggested it was not preferred at all in those simulations whereas Site 2 and 3 were both occupied all the time (Figure S5). Apparently, all of these simulations gave consistent answers, likely driven by the details of the model and force field used. Thus, even though we obtained a different water network in simulations compared to the crystal structure, we still consider this as a success for the tested techniques. In fact, when we checked the experimental electron density map we found that Site 1 has a weaker peak than that of Site 2 and 3, suggesting the probability of observing a water molecule in this site perhaps ought to be lower (Figure S6). However, crystallographic water molecules are typically deposited at 100% occupancy even when density is relatively weak (as is the case for this water) out of a desire to avoid overfitting, complicating interpretation. It is also notable that there were two copies of the protein in the asymmetric unit in the crystal structure with this PDB code (3RLQ) and we used the first chain to prepare our simulations. But in the second chain, water molecules were deposited in both Site 2 and 3 but not Site 1 (unlike in the first chain), further suggesting the uncertainty of the occupancy of Site 1 in the crystal structure – in particular, if we had chosen to compare with this second copy, we would have concluded that Site 1 ought not to be occupied. Thus, our simulation results here seem somewhat consistent with the relatively lower experimental electron density for this water, though we are skeptical that this particular site is favorable at all with the present force field. This analysis also suggests that crystallographic water molecules ought to be more carefully analyzed rather than treating them as simply “there” or “not there”, as some previous studies have done.

**Figure 8:**
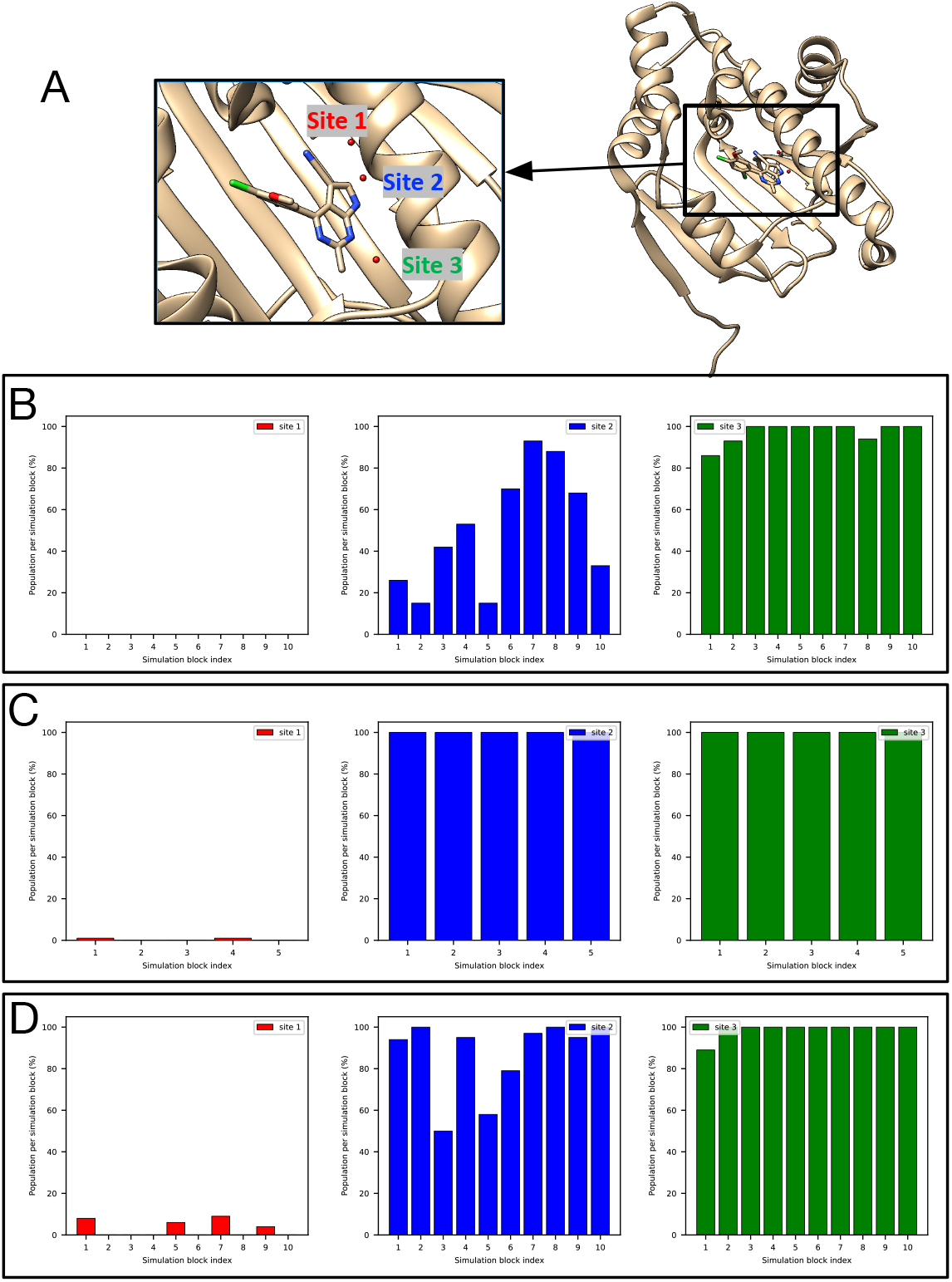
None of our simulations could rehydrate Site 1 (red), though they could rehydrate Sites 2-3 (blue, green) in (A) The HSP90 system (PDB: 3RLQ). Bar graphs show the water occupancy of target sites in a single (B) BLUES simulation (with ordered water molecules removed prior to simulation), (C) *grand* simulation, and (D) unbiased MD simulation.

### 4.4 Lessons we learned about the systems studied

In the following sections, we will discuss what we learned about the systems we simulated, including insights beyond simple analysis of water sampling. We hope these results will aid future work on these systems.

#### 4.4.1 HSP90 (PDB: 2XAB)

As defined above, the reference occupancy is the converged water occupancy from simulations and is used to determine whether we consider a given trial a success. In this case the reference occupancies for all three target sites are 100% as suggested by different simulation techniques (Figure S7). Both BLUES and *grand* can rehydrate three target sites in HSP90 with this ligand (Figure 9). But unbiased MD cannot rehydrate any of the three sites even with much longer simulation times (700 ns). All three sites were highly favorable and none of them could be removed whether simulations started with or without ordered waters. The calculated electron density map agrees well with the experimental electron density map (Figure 9B).

**Figure 9:**
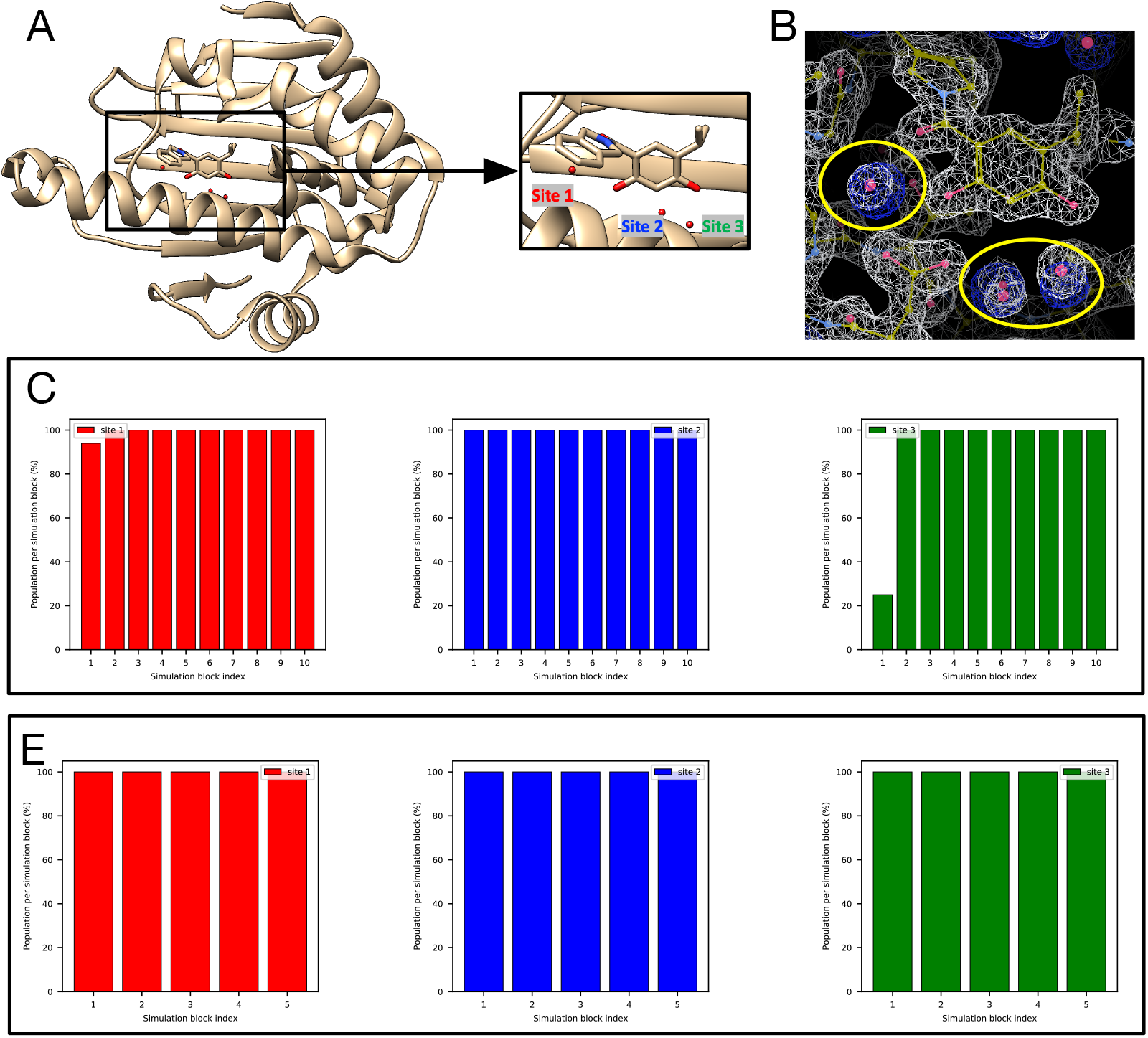
Both BLUES and *grand* simulations can rehydrate all three target sites (Site 1: red, Site 2: blue, Site 3: green) in (A) the HSP90 system (PDB: 2XAB). (B) The calculated electron density map (blue) overlaps with the experimental electron density (2*F_o_ – F_c_*) map (white). The target hydration sites are circled. The calculation is based on a BLUES simulation trajectory where all target sites have occupancies of 100%. Bar graphs show the water occupancy of target sites in a single (C) BLUES simulation, (D) *grand* simulation.

#### 4.4.2 HSP90 (PDB: 2XJG)

Relative to the HSP90 system just prior, the ligand in this case is modified in a way which displaces two water molecules in the binding site. The reference occupancy of the only target site is 100% (Figure S1A-B). MD/BLUES/*grand* can all rehydrate the only target site (Site 1 in Figure 10A) although only one replicate MD simulation could achieve this and it took much longer (280 ns in total, 145.5 million force evaluations, Table 2) than BLUES and *grand* simulations.

**Figure 10:**
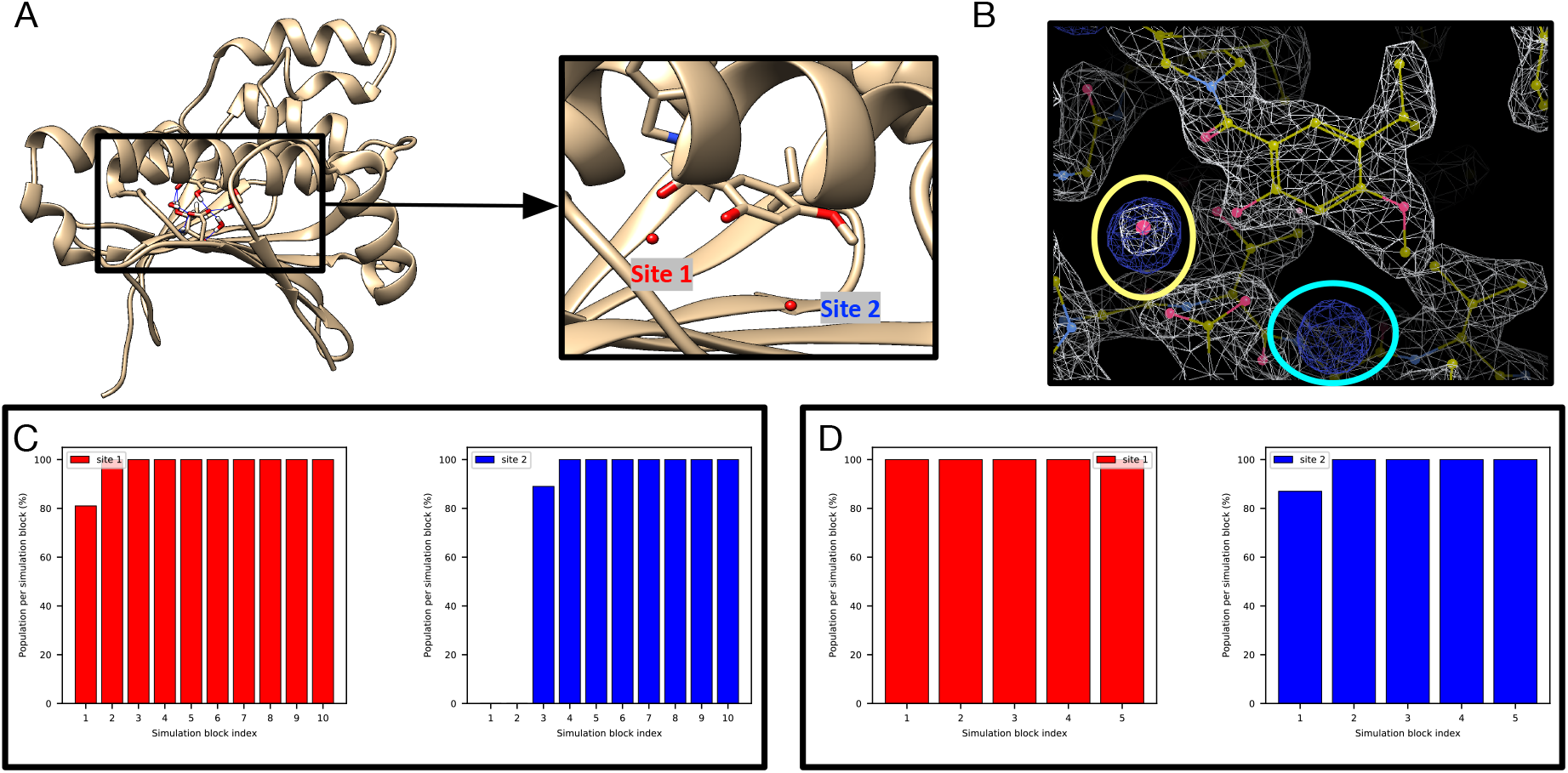
Both BLUES and *grand* simulations can rehydrate the target site (red) in (A) The HSP90 system (PDB: 2XJG). Our results also suggest another favorable site near the binding site in which no waters are deposited in the crystal structure. (B) The calculated electron density map (blue) overlaps with the experimental electron density (2*F_o_ – F_c_*) map (white). The target hydration site is circled in yellow and the extra site is circled in cyan. The calculation is based on a BLUES simulation trajectory. Bar graphs show the water occupancy of target sites in a single (C) BLUES simulation, and (D) *grand* simulation. The number of force evaluations of each simulation block in (C-D) is 1.4 million.

Besides the target site, we found another favorable site (Site 2 in Figure 10A) near the binding site in BLUES/MD/*grand* simulations (Figure 10C-D). The electron density map from our simulations also confirms the existence of this extra water site (circled in cyan in Figure 10B). By checking snapshots extracted from the simulations, we found this water molecule forms a hydrogen bonding network that also involves SER52, ASP93, THR184 and the crystallographic water in Site 1. A previous study of this system also observes this site being occupied in their simulations but no ordered water is deposited in the crystal structure. ^30^ We did not see significant experimental electron density in this water site either. Both that work and our work used same solvent model (TIP3P) and force field for protein (AMBER ff14SB), suggesting this is a force field issue.

#### 4.4.3 HSP90 (PDB: 3RLP)

This additional HSP90 case focuses on a different ligand series (Figure 1) from those above (PDBs: 2XAB, 2XJG). This system has four target water sites (Figure 11A). BLUES simulations (with all ordered water molecules retained prior to simulations) suggest ~100% occupancies for all target sites (Figure 11B).

**Figure 11:**
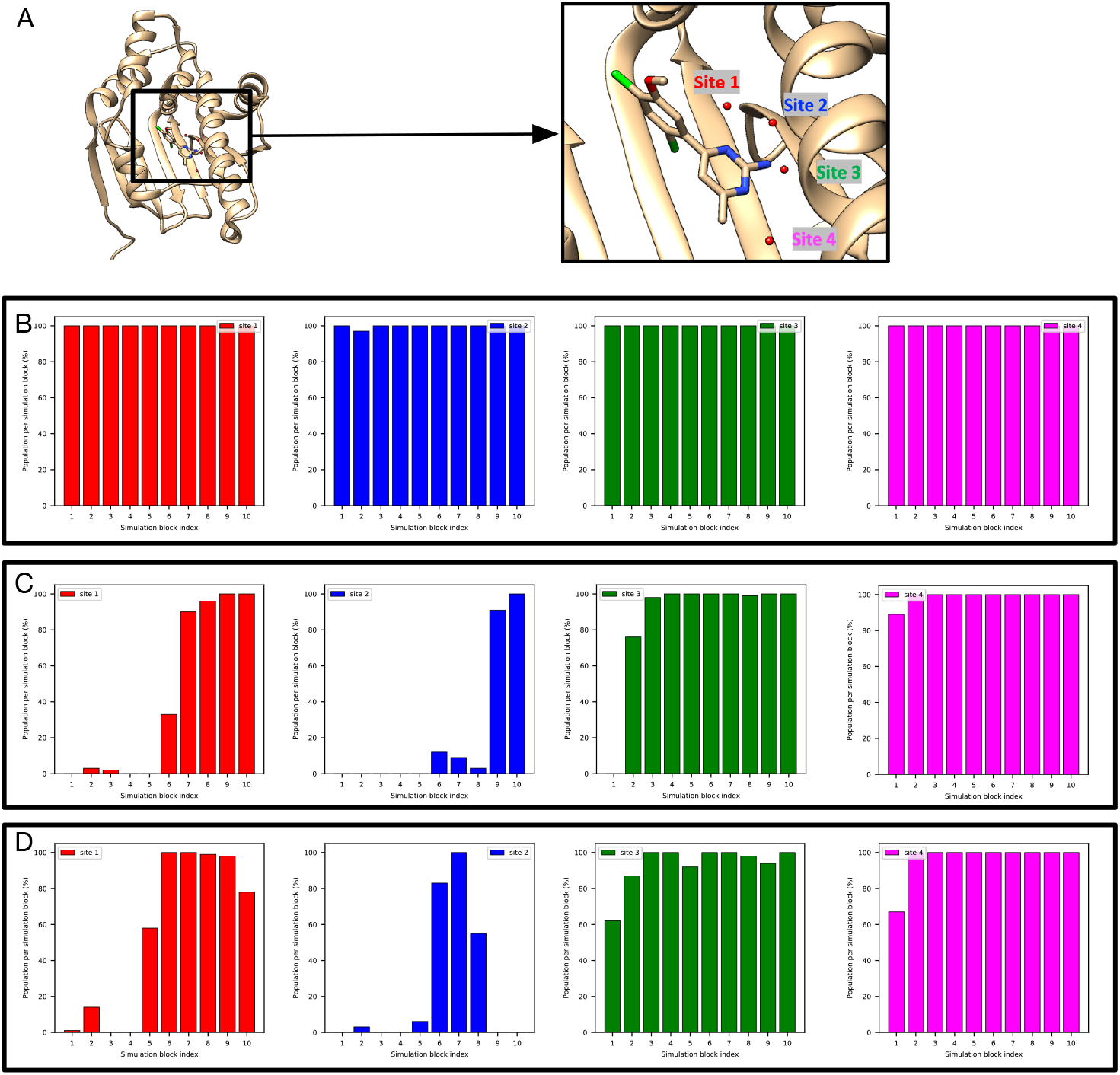
It is more challenging to rehydrate Site 1-2 (red, blue) than Site 3-4 (green, magenta) in BLUES simulations of (A) the HSP90 system (PDB: 3RLP). Bar graphs show the water occupancy of target sites in a single (B) BLUES simulation (all ordered water molecules were retained prior to simulations), (C-D) BLUES simulations (all ordered water molecules were removed prior to simulations).

We found it is more challenging to insert a water molecule in to Site 1 and Site 2 than the other sites in BLUES simulations as it takes longer simulation time to do so (Figure 11C-D). These two sites are also challenging to rehydrate in *grand* simulations and we do not get converged results from *grand* simulations (Figure S8A-B, given that occupancies do not agree).

We then analyzed MD simulations to further check occupancy of target sites especially for Site 1 and 2. We found a very low occupancy of Site 1 in MD simulations (Figure S9). The occupancies of Site 2 and 4 also vary between the two replicates.

The occupancies obtained from simulations vary between replicates so we cannot determine reference occupancies for these target sites. We thus decided to use electron density to assess simulation performance. In this work, when we need to use electron density to compare different simulations, we always compare them using the same analysis method. That said, we do not use success defined based on clustering-based analysis for one simulation technique and compare with other simulations from electron density map analysis. Besides this HSP90 system, we also performed electron density analysis for success and efficiency check for several other systems. The rule we mentioned above applies to all of these systems. The results are shown in Figure S10, S11, S12. We can see all these simulations can reproduce the experimental densities although only one replicate of MD simulations can achieve it. We showed the simulation block when we first saw the success from simulation in Figure 12. In the only successful trial of MD simulation, the peak of Site 1 is a little weak compared to BLUES and *grand* simulations. It is also interesting that MD simulation is more efficient than BLUES simulation in this case (Table 2, Figure 12). The fact that only one trial can rehydrate all target sites highlights the challenges of this system in MD simulations.

**Figure 12:**
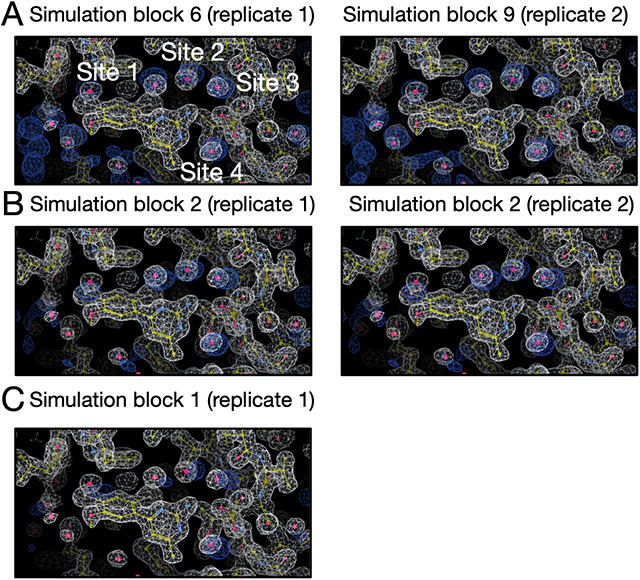
Calculated electron densities (blue) from simulations compared to the experimental densities (white) of the HSP90 system (PDB: 3RLP). Target sites are labeled. We only show the simulation block in which we first observe a good agreement between the calculated and experimental density. All simulations can reproduce the experimental electron densities for target sites but MD simulations can only achieve it in one replicate. Among all simulation techniques, BLUES is the most expensive one to achieve success in this case. Panel A, B and C are results from BLUES, *grand* and MD simulations, respectively. The left and right columns are results from two replicates for each simulation technique.

Besides exploring water sampling issues, we also learned about the protonation state of the ligand. Based on pKa estimates from Chemicalize (a ChemAxon product, https://www.chemaxon.com), there are two possible protonation states for the ligand at the experimental conditions (pH=4.3) (Figure S13A-C). However, the ligand is not stable in the binding site with one of the protonation states and escaped quickly in both unbiased MD and *grand* simulations even at a timescale shorter than 2.5 ns (Figure S13D-E). We didn’t observe such unbinding events in BLUES simulation because the ligand was restrained. The other protonation state of the ligand showed much better stability in the simulations (as long as 700 ns of unbiased MD). Thus, we believe for this system, the ligand protonation state as shown in Figure S13C dominates when the ligand is bound.

#### 4.4.4 HSP90 (PDB: 3RLR)

The modified ligand in this system displaced additional two water molecules from the system mentioned above (HSP90, PDB: 3RLQ). The reference occupancy is 100% (Figure S14). The only target site (Figure S15A) can be rehydrated in the BLUES and *grand* simulations (Figure 4.4.4). However, unbiased MD simulations failed to do so even with a much longer timescale and higher cost (350 ns, 175M force evaluations).

#### 4.4.5 TAF1(2) (PDB: 5I29)

There are five hydration sites in the TAF1(2) system (Figure S16A) and all of them are favorable in BLUES simulations where all water molecules were retained prior to simulations (Figure S17).

In Figure 13C-D we can see all five sites are successfully rehydrated in BLUES and *grand* simulations. In fact, all five water sites were already rehydrated after equilibration simulations (Figure S18). It was surprising to see all five sites rehydrated during the preparation for BLUES simulations given the fact that no biased sampling was applied in equilibration – in other words, these sites were rehydrated while equilibrating with standard MD. We thought this might be because the protein and ligand heavy atoms were restrained to the crystallographic pose in equilibration for BLUES so we performed four additional equilibration simulations (NVT+NPT, see METHODS) using the same restraints as in the original equilibration simulations. With the original equilibration simulation, the five equilibration simulations all return high occupancies for these five target sites (Figure S19). The average occupancy of Sites 1-5 is 73%, 81%, 100%, 83%, 92%, respectively. This is the only system in this work we found that using restraints to maintain the crystal pose in MD simulations improves the probability of successful insertion of water molecules to these target sites.

**Figure 13:**
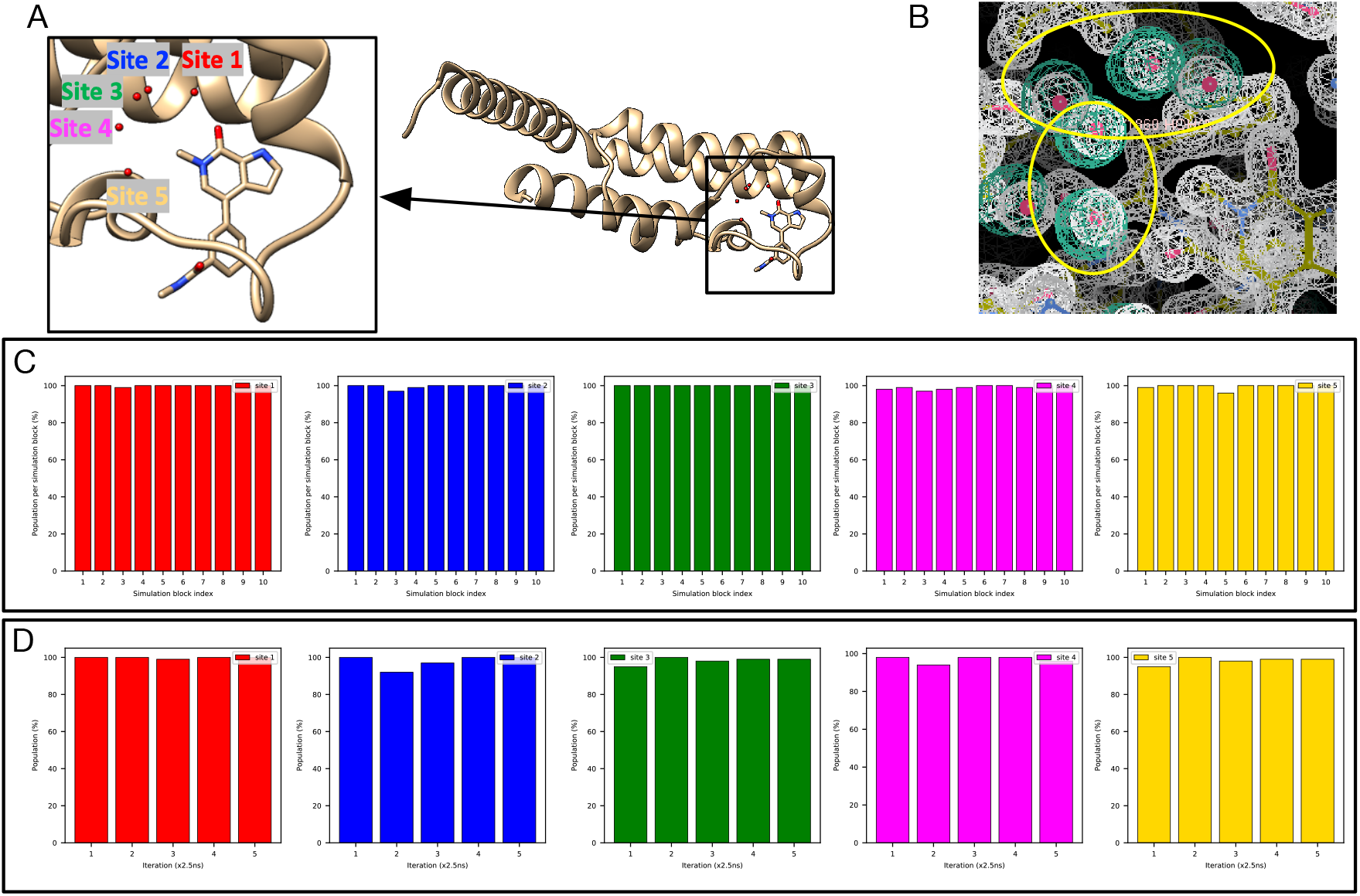
Both BLUES and *grand* simulations suggest high occupancies (close to 100%) for Site 1-5 (red, blue, green, magenta, yellow) in (A) the TAF1(2) system (PDB: 5I29). (B) The calculated electron density map (blue) overlaps with the experimental electron density (2*F_o_ – F_c_*) map (white). The target hydration sites are circled. The calculation is based on a BLUES simulation trajectory where all target sites have occupancies of 100%. Bar graphs show the water occupancy of target sites in a single (C) BLUES simulation and (D) *grand* simulation.

In this system, we also found both BLUES and *grand* simulations could remove the water molecules from the sites after they were occupied and then rehydrate them again, indicating we could converge population estimates. This ability to sample multiple water transitions into and out of the sites is not common for the systems studied here. One possible reason for the additional ease of sampling here could be that this binding site is large and more exposed to the bulk solvent than binding sites in the other systems examined.

In MD simulations, however, we found occupancies of these target sites were lower than those in BLUES and *grand* simulations especially for Site 3 and 4 (Figure S20). One possible reason could be the lack of position restraints (compared to BLUES simulations) and much longer timescale (compared to *grand* simulations) in MD simulations. However, our clustering-based analysis may not reveal the true occupancies of these sites. We noticed the ligand is flexible in the binding site in simulations and this binding site is more exposed to the bulk solvent. So water sites change locations as the protein and ligand rearrange. In our clustering-based analysis, we found many sampled sites from simulations and this posed challenges in determining which of the target sites these corresponded to (Figure S21). That said, we may miss some sites in our analysis which may lead to the lower occupancies we observed in this case.

As noted above, electron density based analysis can be very useful in this case as we do not focus on single sites. Instead, we are comparing calculated electron densities to the experimental densities; this analysis has a higher tolerance for water movements due to protein/ligand motions. Thus, instead of using reference occupancies, we decide to use electron density maps to check the performance of different simulation techniques. In fact, this is not the only case where we found our clustering-based analysis was not robust; this occurs in several cases (below).

We calculated the electron density for each simulation block (Figure S22,S23,S24) and we found all simulation techniques can reproduce the experimental densities within the first simulation block based on visual inspection. As noted in Section 3, we used a contour level of 3 sigma for calculated water electron density maps and 1.5 sigma for experimental protein/water maps across all systems. Since each simulation block has a different number of force evaluations for each simulation technique (BLUES: 18 million, MD: 35 million, *grand*: 1.4 million) in igure S22,S23,S24, we performed additional analysis in which each simulation block has the same number of force evaluations (1.4 million). The results are shown in Figure 14. Since the longer simulations in Figure S22,S23,S24 already showed a good agreement between the calculated and experimental electron densities, here we only track the simulation block at which we first observe a good agreement between the calculated electron densities and experimental densities in this additional analysis. We can see both BLUES and *grand* simulations can reproduce the experimental densities within simulation block 1 (1.4 million force evaluations, Figure 14A,C) whereas it takes longer for MD to achieve such agreement (simulation block 5 (7 million force evaluations) and 9 (12.6 million force evaluations), Figure 14B), indicating a substantial efficiency gained with BLUES and *grand* simulations compared to normal MD simulations.

**Figure 14:**
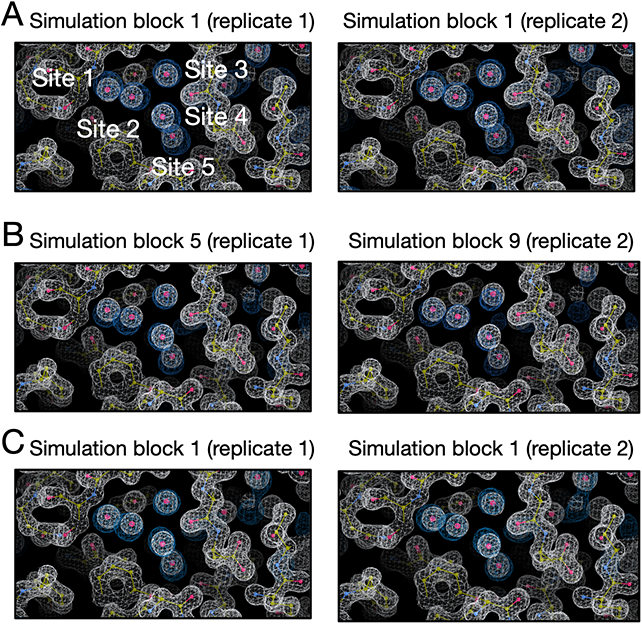
Calculated electron densities (blue) from simulations compared to the experimental densities (white) of the TAF1(2) system (PDB: 5I29). Target sites are labeled. We only show the simulation block when we first observe a good agreement between the calculated and experimental density. All simulations can reproduce the experimental electron densities for target sites but MD simulations are more expensive to achieve it. Panel A, B and C are results from BLUES, MD and *grand* simulations, respectively. The left and right columns are results from two replicates for each simulation technique. The number of force evaluation for each simulation block is 1.4 million in this analysis.

#### 4.4.6 TAF1(2) (PDB: 5I1Q)

A modification of the ligand in the TAF1(2) system discussed above changes the water network in the binding site (Figure 15A). BLUES simulations return converged occupancies for all target sites (Figure 15). We can see Site 5 has a lower occupancy than other 4 sites and we observed more transitions in this site. We found the location of this site is more exposed to bulk solvent. So it is likely to have more transitions between this site and bulk solvent compared to the other 4 sites which are more buried.

**Figure 15:**
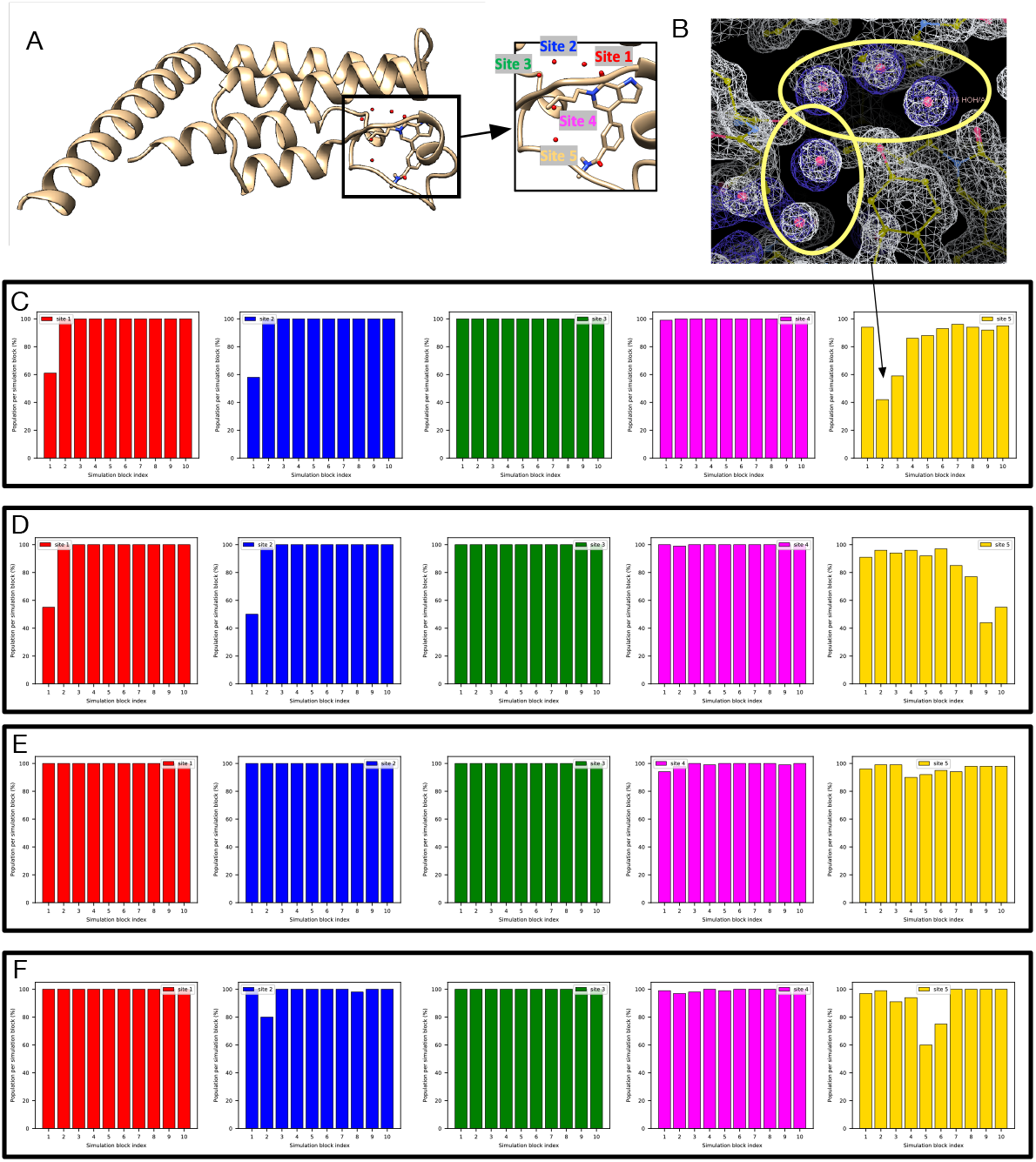
BLUES simulations return converged occupancies for all target sites (Site 1: red, Site 2: blue, Site 3: green, Site 4: magenta, Site 5: yellow) in (A) The TAF1(2) system (PDB: 5I1Q). (B) The calculated electron density map (blue) overlaps with the experimental electron density (2*F_o_ – F_c_*) map (white). The target hydration sites are circled. The calculation is based on a BLUES simulation trajectory shown in (C). Bar graphs show the water occupancy of target sites in BLUES simulations with ordered water molecules (C-D) removed and (E-F) retained prior to simulations.

In MD simulations, with/without ordered water molecules the occupancy of Sites 4 and 5 does not converge (Figure S25). On average MD simulations with all ordered water molecules retained prior to simulations return an occupancy of 50% and 40% for Site 4 and 5, respectively. However, in MD simulations with all ordered water molecules removed prior to simulations, Site 4 and 5 only have an average occupancy of 5% and 14%. The two *grand* simulation replicates do not converge either (Figure S26). In one trial, Site 1-3 are almost 100% occupied but the occupancies of these three sites are much lower in another trial. This is due to the issue of the ligand motion discussed above. We also found both *grand* trials return a lower occupancy for Site 4 and 5 and they do not converge. In trial 1, Site 4 has an average occupancy of 46% whereas in trial 2 it is only 15%. Site 5 has an average occupancy of 44% and 59% in trial 1 and 2, respectively. We also tried to extend the simulation timescale (25 ns) by a factor of two (from 12.5 ns). However, the results were still not converged.

Similar to another TAF1(2) system above, the clustering-based analysis does not return a clear water network due to the flexibility of the protein and ligand (Figure S21). So it is unclear whether the simulations are not converged or our clustering-based analysis does not accurately reflect the occupancies of these sites from simulations. Because of this, we switched to electron density analysis since we found it more robust in analyzing simulation data for another TAF1(2) system where we encountered similar issues.

In Figure S27 we can see BLUES simulations achieve a good agreement between the calculated electron densities and experimental densities in both replicates within the first simulation block. Both replicates of MD simulations failed to rehydrate all target sites (Figure S28). One replicate of *grand* simulation achieves success (Figure S29A), simulation block 6) but the other one fails (Figure S29B). Since each simulation block has different number of force evaluations for each simulation technique (BLUES: 18 million, *grand*: 1.4 million) in Figure S27 and S29, we performed additional analysis in which each simulation block has the same number of force evaluations (1.4 million). The results are shown in Figure 16. We only show the simulation block where we first observe a good agreement between the calculated electron densities and experimental densities. Both BLUES and *grand* simulations have similar efficiency in this case as rehydrating all target sites takes 8.4 million and 7 million force evaluations for BLUES replicates and 8.4 million force evaluations for the only successful *grand* replicate (Figure 16). But if we include force evaluations in equilibration phase, then *grand* simulations are more efficient in this system (Table 1).

**Figure 16:**
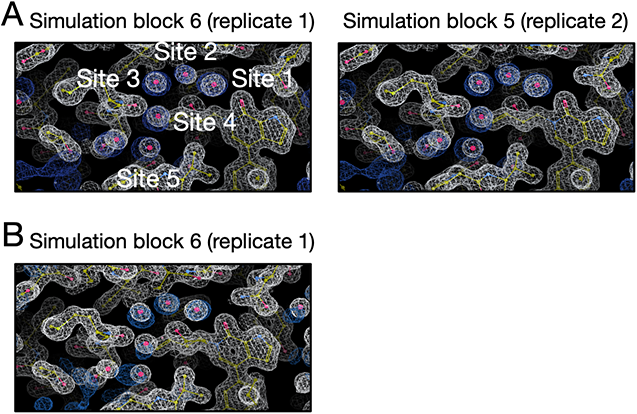
Calculated electron densities (blue) from simulations compared to the experimental densities (white) of the TAF1(2) system (PDB: 5I1Q). Target sites are labeled. We only show the simulation block when we first observe a good agreement between the calculated and experimental density. Both BLUES replicates can reproduce the experimental electron densities for target sites but only one replicate of *grand* simulations can achieve this. Panel A and B are results from BLUES and *grand* simulations, respectively. The left and right columns are results from two replicates for each simulation technique. The number of force evaluation for each simulation block is 1.4 million in this analysis.

We found Sites 4 and 5 are especially challenging for MD simulations and *grand* simulations. We also saw the occupancy of Site 2 substantially between the two replicates of *grand* simulations (Figure S29B). As we mentioned above, this is due to ligand motion that blocks the successful insertion of water molecules (Figure 4).

#### 4.4.7 BTK (PDB: 4ZLZ)

This BTK system has one target hydration site in the binding site (Figure S30A), bridging the protein and ligand as shown in prior work.^77^ MD and BLUES simulations do not converge to the same occupancy for the target site (Figure S30). In BLUES simulations, the target site is 100% occupied whereas in MD simulations it is occupied only 60% (averaging over all simulation blocks in Figure S30C-D). We have seen similar discrepancies between MD simulations and BLUES simulations in those TAF1(2) systems. This is likely due to the use of position restraints on heavy atoms in BLUES simulations so that the protein-ligand complex system always maintains the crystal conformation. Such restraints were not used in MD simulations and both the protein and ligand were more flexible than those in BLUES simulation. We noticed fluctuations in MD simulations (Figure S30C-D) between blocks and replicates. This suggests the simulations are not converged yet. Meanwhile, we also found the protein and ligand are flexible in simulations and our clustering-based analysis returns many sampled sites (similar to Figure S21) just like we observed in two TAF(1)2 systems. So the occupancies from MD simulations are not reliable since we may miss some sites in our analysis. Thus we decided to use electron density analysis in this case since it works well when occupancies from clustering-based analysis are questionable (e.g., TAF1(2) systems).

In Figure S31, S32, S33, we can see that all simulations rehydrate the target site within the first simulation block. Since each simulation block has different number of force evaluations for each simulation technique (BLUES: 18 million, MD: 35 million, *grand*: 1.4 million) in Figure S31, S32, S33, we performed additional analysis (Figure 17)so that each simulation block has the same number of force evaluations (1.4 million). We can see both replicates of *grand* simulations can rehydrate the site within the first simulation block. BLUES and MD both have one replicate that can achieve a good agreement between the calculated and experimental electron density within the first simulation block while the other replicate takes longer to achieve it (2.8 million force evaluations for BLUES; 8.4 million force evaluations for MD).

**Figure 17:**
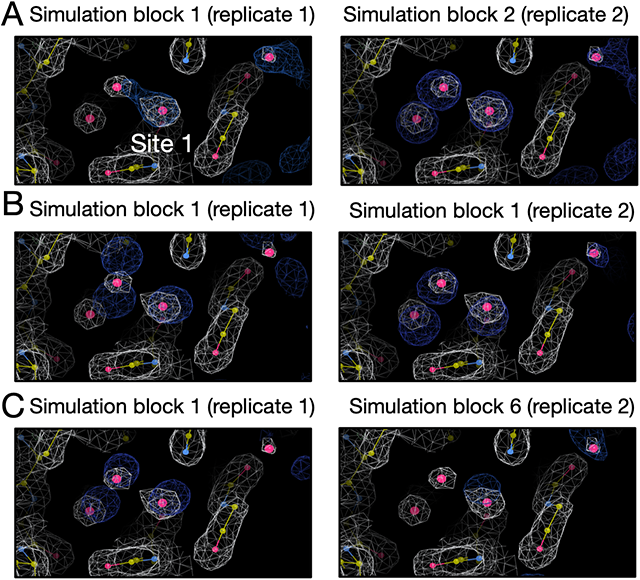
Calculated electron densities (blue) from simulations compared to the experimental densities (white) of the BTK system (PDB: 4ZLZ). The target site is labeled. We only show the simulation block when we first observe a good agreement between the calculated and experimental density. All simulations can reproduce the experimental electron densities. Panel A, B and C are results from BLUES, *grand* and MD simulations, respectively. The left and right columns are results from two replicates for each simulation technique. The number of force evaluation for each simulation block is 1.4 million in this analysis.

#### 4.4.8 Thrombin (PDB: 2ZFF)

We already discussed challenges in rehydrating the target site in the thrombin system in Section 4.3.2. Other than water sampling issues, two possible protonation states of the ligand are suggested based on the pKa calculations using Chemicalize (ChemAxon, https://www.chemaxon.com) at pH = 7.5 (Figure S34). In our simulations, both protonation states were stable in the binding site when no restraints were applied, meaning that we cannot tell from this data which is preferred or dominant. This is different from the case of HSP90 (PDB: 3RLP) in which only one ligand protonation state shows reasonable stability of the ligand whereas the other one leads to unbinding of the ligand very quickly.

## 5 DISCUSSION

Although it is well known that water molecules can influence different biological processes (e.g., protein-ligand binding)^6 –^ and computation is frequently used to explore such processes, systematic comparisons of water sampling techniques are infrequent. However, we believe such comparisons are important since we can only improve the methods after we learn where and how they fail.

In this work, we studied the sampling of buried waters in binding sites using several different simulation techniques. We studied a range of protein-ligand systems, most of which have hydration sites which vary their occupancy as different congeneric ligands bind.

One important lesson we learned from this work is that neither clustering-based analysis nor electron density map analysis alone can adequately capture a complete picture of water occupancy and rearrangement in the full range of outcomes we encountered in our simulations. The use of clustering-based analysis provides coordinates of representative water sites occupied by favorable water molecules in simulations which can be compared to the experimental crystal structures. In this case, occupancy information can also be obtained by calculating the frequency of favourable regions being occupied by water molecules in simulations, enabling more robust quantitative analysis.

This clustering-based approach compares the results from simulations with crystallographic water molecules which are deposited by crystallographers based on the electron density maps. However, those crystallographic waters are based on interpretations of the underlying data, introducing the potential for human bias and/or errors.^85, 86^ Additionally, crystallographic water molecules are typically deposited at 100% occupancy even if the experimental density is relatively weak and might suggest lower occupancy. This poses difficulties in directly comparing between simulation-predicted and experimental occupancies. Water occupancies are typically not refined in order to avoid overfitting, but still this limitation precludes direct comparison between simulations and experiments. This, however, is a limitation which cannot be addressed within the scope of the present work. Still it is important to keep in mind that the water molecules in the crystal structure are not always reliable, and thus differences between water network shown in the crystal structure and revealed in the simulation do not necessarily lead to the conclusion that something is wrong in the simulation.

Additionally, the clustering-based analysis works best when the protein/ligand are restrained or stable in simulations. When they are not restrained or when they are highly flexible, as we observed in this work, protein/ligand motions may interfere with the water network from clustering analysis as water sites change locations as the protein and ligand rearrange. Correspondingly, simulations discover many water sites, making it difficult to compare with crystallographic waters since it is not easy to assign water sites populated from simulations to the crystallographic sites for comparison (Figure S21), though possibly this issue could be overcome by advances in analysis.

An alternative method is comparing calculated electron density of water molecules with the experimental electron density maps (2F*_o_*-F*_c_*). This analysis brings us one step closer to the original experimental data than does analyzing discrete water molecules in the provided structure deposited in the PDB. We find that this approach also helps with analysis of our simulation, since we are able to compare regions of significant water occupancy rather than limit our analysis to a single site with specific coordinates. Especially when the clustering-based analysis failed in providing reliable occupancies of target sites in the TAF1(2) and BTK system in this work, we found using electron density map analysis was more robust to assess the performance of simulation techniques. However, compared to the clustering-based analysis, it requires extra work to calculate occupancies of water sites.

Although not shown in this work, another potential advantage of using electron density maps in the analysis is that doing so offers an opportunity to compare with the F*_o_*-F*_c_* map (difference map) so that differences between simulated and deposited water network in the crystal structure can be further analyzed. As we mentioned earlier, both force fields and crystal structures are not perfect so it is not surprising to see water sites populated in simulations differ from those in crystal structures. Comparing the calculated electron density map with the difference map from experimental densities could help to assess simulation performance in recovering all hydration sites in the crystal structure. For example, if the water sites sampled in the simulation are in the region with positive peaks (shown as green in electron density maps), it is possible that the simulation captures the water molecules that are suggested by experimental electron density but have not been modelled by crystallographers. In contrast, if the water sites suggested by simulations are in a region with no peaks then it is possible that the force field is not accurate and placed water molecules in sites which should be devoid of water. It is notable that the complete interpretation of difference map can be complicated and also relates to factors other than water molecules (e.g., ions, protonation states, co-solvents, etc.) but in any case, these maps provide information for additional consideration when examining discrepancies between simulations and crystal structures.

Based on our experience in this work, we suggest researchers use both approaches in the study of water sites, to allow the analysis of both specific, discrete, well-defined water sites and broader favorable regions that are occupied by water (sometimes even sporadically) in simulations. In addition, applying two approaches allows for cross validation to ensure consistency.

The analysis we performed in this work did not consider potential biases introduced by using position restraints on the protein and ligand and compared the results directly with normal MD simulations where no restraints were used. Without restraints, our enhanced sampling methods often simply did not achieve adequate acceptance. Ideally, one would correct for the effect of these restraints such as with reweighting techniques.^87^ However, our restraints here were relatively strong (10 kcal mol*^−^*^1^ Å*^−^*^2^ on all heavy atoms of the protein and all atoms of the ligand). Reweighting techniques with such strong restraints would likely result in a very small number of effective samples contributing to final estimates, and thus introduce substantial statistical uncertainty. Thus, reweighting was not employed here. But in future work with weaker restraints, reweighting techniques may be helpful to correct for the effects of restraints when computing properties like the hydration site occupancy. In a previous study that found restraints were important in recovering crystallographic waters using MD simulations,^66^ a spring constant of 0.5 kcal mol*^−^*^1^ Å*^−^*^2^ was used, as a way of avoiding artificial ordering; ^88^ it would be interesting to explore using this value in further studies.

Another issue making our analysis more difficult is that there is no well-established definition for successful water rehydration/sampling in simulations. The definition we used in this work is reasonable but definitely not the only possible definition. This definition is important since it may affect the assessment of different methods. Depending on the sampling in simulations, a system dependent success criterion may be necessary. In this work, we first tried to use an occupancy-based criterion and it worked well in half of studied systems (5 systems). However, we found in the other 5 systems, we either could not get converged occupancies from our MD reference simulations (all ordered water molecules were retained prior to simulations) or our clustering analysis was not robust in dealing with these systems (i.e., it is difficult to determine which target sites the sampled sites corresponded to, Figure S21). Instead, using an electron density-based criterion was more successful in these such cases. In fact, using electron density-based analysis can also work in those cases where clustering-based analysis works. But as we mentioned, extra work is needed to calculate occupancy information in electron density analysis if such information is desired.

These challenges highlight that the challenging topic of water occupancy still requires more attention, both in terms of computational modeling and experimental interpretation (since crystallographic waters currently seem to be deposited only at full occupancy, even if the underlying density is weak). Based on our experience in this work, inspection of experimental crystal structures provides no clear indication as to which buried waters in the binding sites will be difficult to rehydrate in simulations. For example, the HSP90 system (PDB: 3RLR) only has one water molecule in the binding site but poses challenges for MD simulations. Small modifications in a congeneric series of ligands could lower the chances for successful rehydration. Even with the same receptor (e.g., HSP90), the difficulty of rehydrating all target sites in the binding sites varies between ligands with minor structural differences (Table 2). On the experimental side, it would be more helpful if crystallographers would deposit more information on water molecules in crystal structures, such as including water occupancies in the refined model. We hope this work will draw more attention to these water-related issues so that we can improve our understanding of roles of water molecules in the active sites of the protein targets in future work.

None of the methods we studied in this work can handle water sampling perfectly although *grand* appears more robust than MD and BLUES. Even using GCMC in equilibration phase is helpful for adequate water sampling of target sites in several systems (Table 2). However, we also observed that protein/ligand motions may impair *grand* performance in water rehydration. These motions are expected in simulations when no restraints were applied to the protein/ligand but may take timescales beyond the typical free energy calculation simulation time in a single trial (e.g., *>* 50 ns). Restraining the protein/ligand may avoid this issue but results of this approach are highly system dependent. Alternatively, applying restraints only in the equilibration runs and removing them in the production runs is also a way to alleviate this issue. Either way requires prior knowledge of the simulated structure with target hydration sites occupied (e.g., crystal structures, docked poses or homology models) which may not be always available in blind challenges or in a discovery setting. Moreover, our results on one thrombin system (PDB: 2ZFF) suggest protein/ligand flexibility is sometimes necessary for successful water rehydration attempts (Figure 6) to allow response to water insertion. Unfortunately, such information may not be available in advance, impairing *grand*’s predictive power.

BLUES enhances water sampling relative to normal MD but appears less efficient than *grand* (Table 1, 2). In the BLUES protocol used in this work, we deployed 3000 iterations of NCMC moves in a single simulation block, accumulating 18 million force evaluations (including both MD and NCMC steps) which is equivalent to 12 ns simulation time. In *grand*, a typical single run (1.4 million force evaluations, 2.5 ns) performs 125000 GCMC moves in which each GCMC move attempts to insert/remove a water molecule in to the site. This is about 42 times more attempts than BLUES (3000 attempts) in a single run in this work. The difference between the protocols of BLUES and *grand* in this work is due to the fact that *grand* performs instantaneous water insertion/deletion through GCMC moves but BLUES alchemically turns off/on the interactions of the water molecule with its surrounding environment before and after translating it to a new location. Thus, for one water insertion attempt, BLUES is more expensive than GCMC which explains the performance differences between BLUES and *grand* (Section 4.2, Table 2). Additionally, *grand* applies GCMC moves during the equilibration phase and can help water sampling in target sites (Table 2) whereas BLUES only runs normal MD.

In theory, BLUES has potential in rehydrating water sites in the binding site where protein sidechain reorientation is required for successful attempts whereas instantaneous insertion of water molecules by *grand* may fail due to atomic clashes. It is notable that current performance of BLUES relies on the use of restraints on the protein/ligand which keeps the protein cavities from the protein cavities from quickly collapsing. But in future work we could extend BLUES to allow more complex moves, such as a combination of sidechain rearrangement and water hopping moves so that there is no need to restrain the whole protein/ligand but only regions where are not part of the binding/hydration target sites. But this is more appropriate when prior knowledge of the system (e.g., binding/hydration site location, sidechain/ligand motions) is available.

Normal MD simulations encountered difficulties in rehydrating each target site in most of the systems studied here. Even in those cases which were successful, MD simulations were more expensive than *grand* and BLUES. In addition, whereas restraints yielded improved rehydration performance for *grand* and BLUES in most systems, using restraints in MD simulations only improved rehydration in a single system. The failure of restraints to improve MD rehydration here might be due to the competing benefits of increased fidelity of the hydration-site structure vs. the kinetic barriers to rehydration introduced by increasing the stiffness of the protein, when protein atoms must move to create a path to the hydration site. Such an explanation is consistent with the previous finding that using harmonic restraints improved the ability of MD simulations to recover the average crystallographic water structure,^66^ when waters are not removed after the solvation step. It is also possible that using a smaller spring constant than the present one of 10 kcal mol*^−^*^1^ Å*^−^*^2^ would improve rehydration in the case of normal MD (the previous study used a spring constant of 0.5 kcal mol*^−^*^1^ Å*^−^*^2^ study^66^), by lowering barriers to protein/ligand rearrangements that are needed for inserting water molecules.

In five systems studied here, the hydration sites stayed occupied (100%) in the simulation after the water was successfully inserted, suggesting these sites are highly favorable with the force field. Ideally, we would obtain water site occupancy estimates from simulations with reversible transitions of water molecules into and out of such sites. However, for highly favorable hydration sites, such transitions were not observed in either BLUES or *grand* simulations. One way to solve this issue could be to perform more selective move proposals so that more sampling can be focused on selected regions (e.g., target water sites) instead of a broadly defined spherical region as it is in the current BLUES settings. One way to test this idea is to combine the latest move type in BLUES, molecular darting moves ^57^ (moldarting), with current water hopping moves. That is, we can identify regions where water is favorable in simulations. Then, we can use moldarting to propose NCMC moves between these regions for enhanced sampling to obtain more reliable estimate of hydration site populations.

## 6 CONCLUSION

In this work we assessed MD/BLUES/*grand* performance in water sampling using a range of protein-ligand systems. Our results suggest both BLUES and *grand* enhance water sampling relative to normal MD, and *grand* is more robust than BLUES. The lessons we learned about these methods may help the broader community and point to further opportunities for improvement. We also discussed what we learned about each system studied in this work and hopefully these insights are useful for future work on these systems. We also highlighted issues in analyzing water sampling, and we hope that this work will draw more attention to this topic.

## 7 ACKNOWLEDGEMENTS

D.L.M. appreciates financial support from the National Institutes of Health (R01GM108889 and R01GM132386). DLM and YG also appreciate financial support from XtalPi. We appreciate the Open Force Field Consortium for its support of the Open Force Field Initiative, which provided software infrastructure used in this work. M.E.W. was supported by the Exascale Computing Project (No. 17-SC-20-SC), a collaborative effort of the U.S. Department of Energy Office of Science and the National Nuclear Security Administration, and the University of California Laboratory Fees Research Program (No. LFR-17-476732). The author also appreciates the insightful discussions of prior work with Ido Ben-Shalom, Gregory A. Ross, Matteo Aldeghi and Oliver J. Melling for previous work done on some of the systems studied in this work and suggestions on the analysis. The author thanks help of BLUES package usage and simulation preparation from Samuel Gill, Teresa Danielle Bergazin, Ĺea El Khoury and Jeffrey Wagner.

## 8 ASSOCIATED CONTENT

### Supporting Information Available

Supporting information is available free of charge via the Internet at http://pubs.acs.org.

Supporting tables of parameters used in *grand* simulations; supporting figures of water occupancy of target sites and experimental/calculated electron density maps.

Input files for simulations and scripts for analysis are freely available at https://github.com/MobleyLab/water_benchmark_paper.

Simulations were performed using the open-source package BLUES (v0.2.4, https://github.com/MobleyLab/blues), OpenMM (v7.4.2, https://github.com/openmm/openmm), and *grand* (v1.0.0 and v1.0.1, https://github.com/essex-lab/grand).

Analysis was performed using Mdtraj (v1.9.4, https://github.com/mdtraj/mdtraj), *grand* (v1.0.0 and v1.0.1, https://github.com/essex-lab/grand), CCTBX (v2021.1, https://github.com/cctbx/cctbx_project), LUNUS (https://github.com/mewall/lunus), Phenix (v1.91.1, https://www.phenix-online.org), Coot (v0.9.4, installed with Phenix), CCP4 (v7.1, https://www.ccp4.ac.uk).

## 9 Notes

D.L.M. is a member of the Scientific Advisory Board of OpenEye Scientific Software and an Open Science Fellow with Silicon Therapeutics.

## 10 For Table of Contents Only

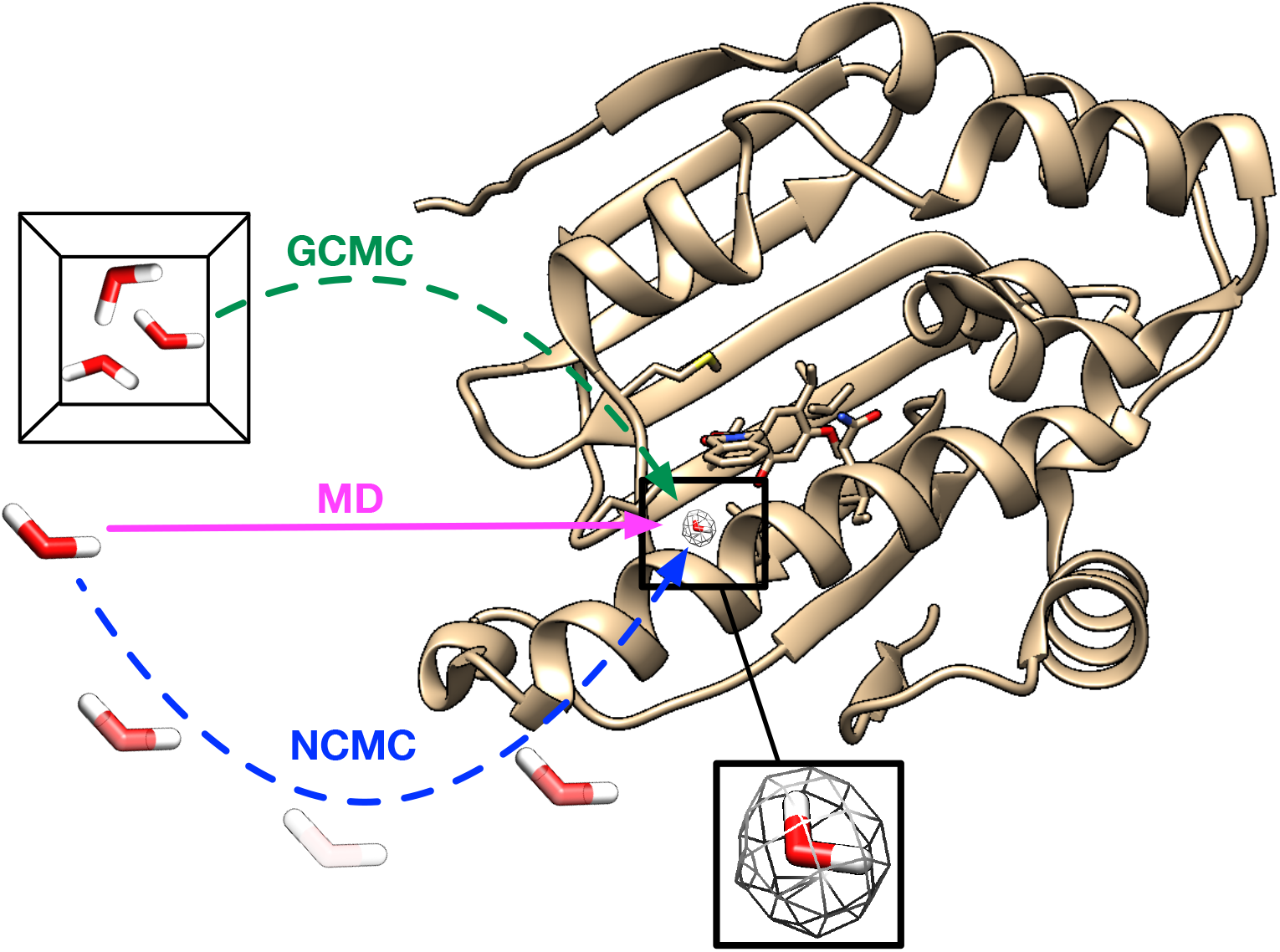

