## Supporting Information for "Enhancing Sampling of Water Rehydration on Ligand Binding: A Comparison of Techniques"

### 11 Supplementary Information

*Corresponding Author

### 12 Supporting Tables

Table S1: The excess chemical potential and standard state volume of water used in *grand* simulations at different temperatures for different systems. Both BLUES and MD simulations were performed at the same temperature for these systems as in *grand* simulations.

| Temperature (K) | Excess chemical potential (kcal/mol) | Standard state volume (Å$^3$) | Systems and PDB IDs |
|---|---|---|---|
| 278 | -6.34 | 29.823 | PTP1B (2QBS)<br>HSP90 (2XAB, 2XJG)<br>Thrombin (2ZFF)<br>BTK (4ZLZ)<br>TAF1(2) (5I1Q, 5I29) |
| 286 | -6.19 | 30.035 | HSP90 (3RLP, 3RLQ, 3RLR) |

### 13 Supporting Figures

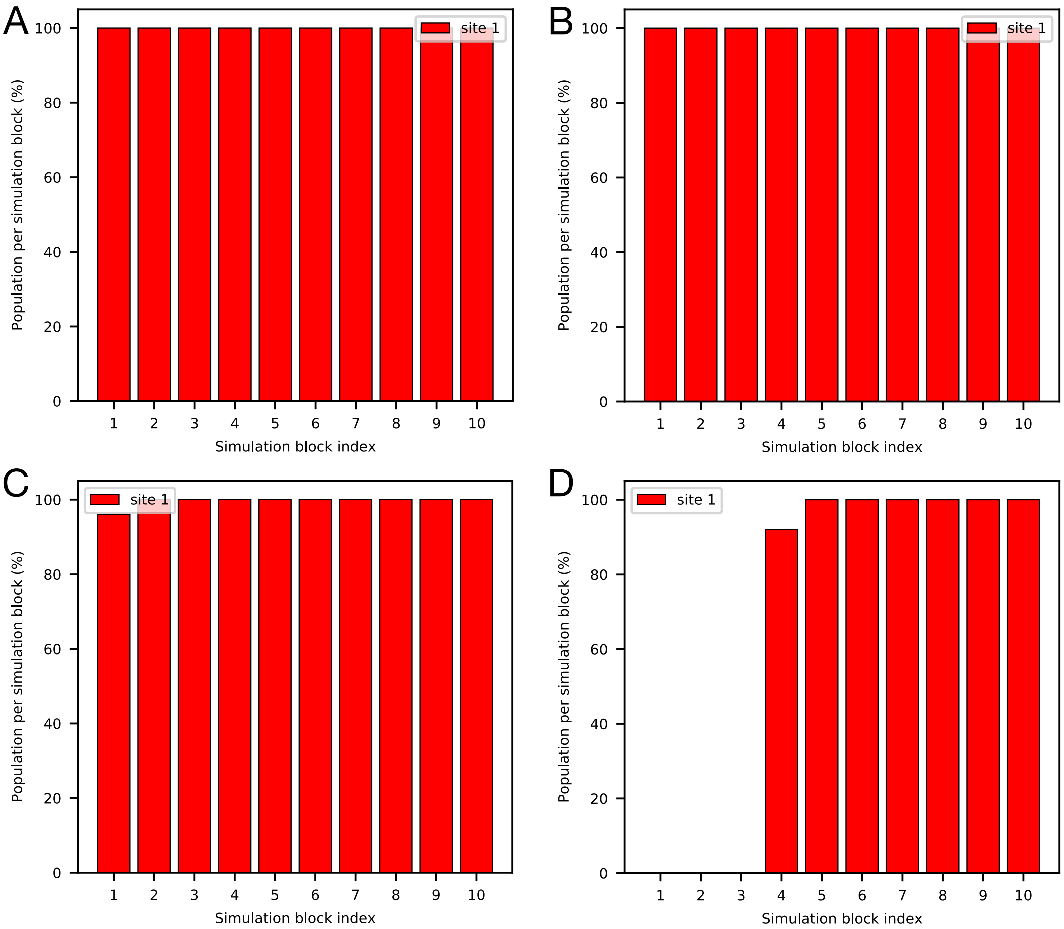


Figure S1: All simulations converge to the same occupancy (100%) of the target site. Bar graphs show the water occupancy of the target site of the HSP90 system (PDB: 2XJG) in (A) BLUES simulation (ordered water retained) (B) MD simulations (ordered water retained) (C) BLUES simulation (ordered water removed) and (D) MD simulation (ordered water removed).

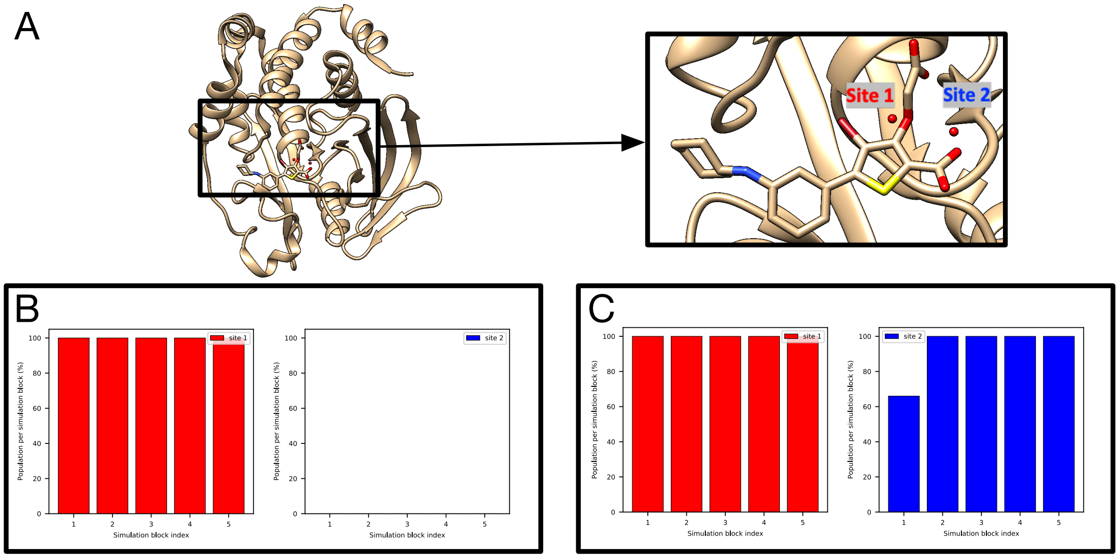


Figure S2: *grand* simulations successfully rehydrate Site 1 in both trials but only are able to rehydrate Site 2 in one trial. (A) The PTP1B system (PDB: 2QBS) and target water sites (Site 1: red, Site 2: blue). (B-C) Bar graphs show the water occupancy of target sites in *grand* simulation (no restraints).

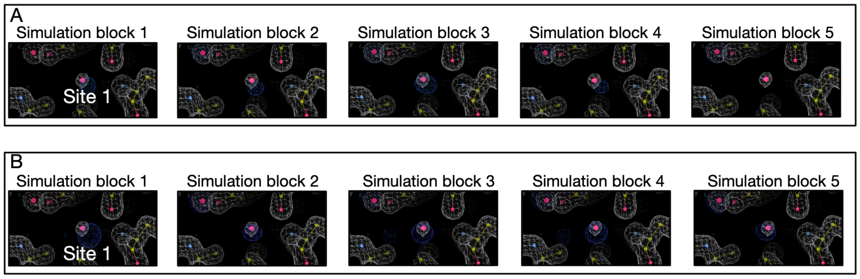


Figure S3: Calculated electron densities (blue) from *grand* simulations agree well with the experimental densities (white) of the thrombin system (PDB: 2ZFF). The target site is labeled. Panel A and B are results from two replicates.

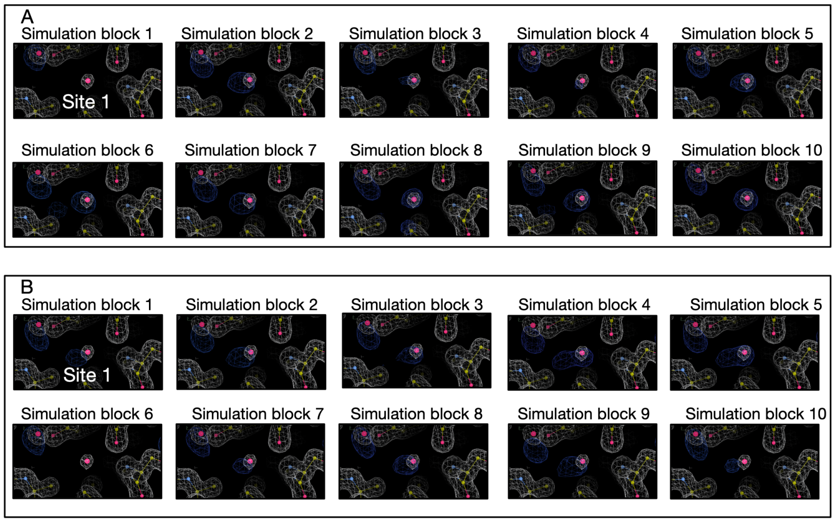


Figure S4: Calculated electron densities (blue) from MD simulations agree well with the experimental densities (white) of the thrombin system (PDB: 2ZFF). The target site is labeled. Panel A and B are results from two replicates.

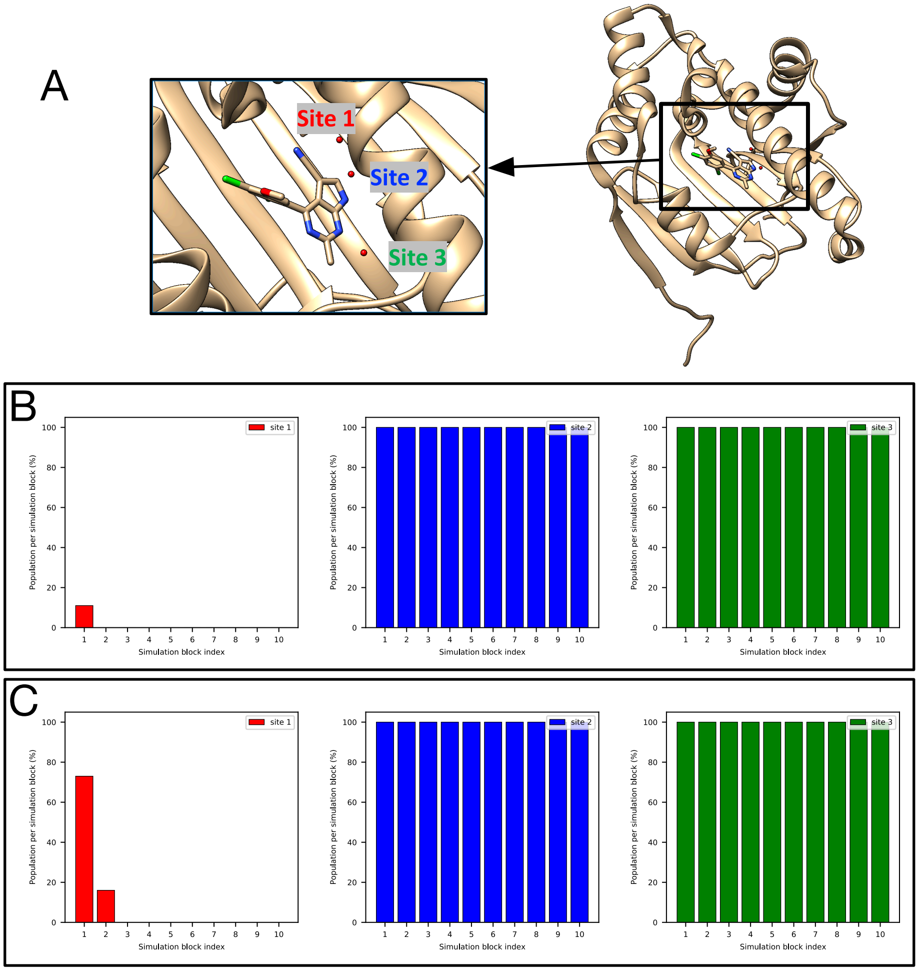


Figure S5: Both BLUES and MD simulations suggest a low occupancy of Site 1 in (A) the HSP90 system (PDB: 3RLQ). Bar graphs show the water occupancy of target sites (Site 1: red, Site 2: blue, Site 3: green). (B) in a single BLUES simulation and (C) unbiased MD simulation. In both simulations, ordered water molecules were retained prior to simulations.

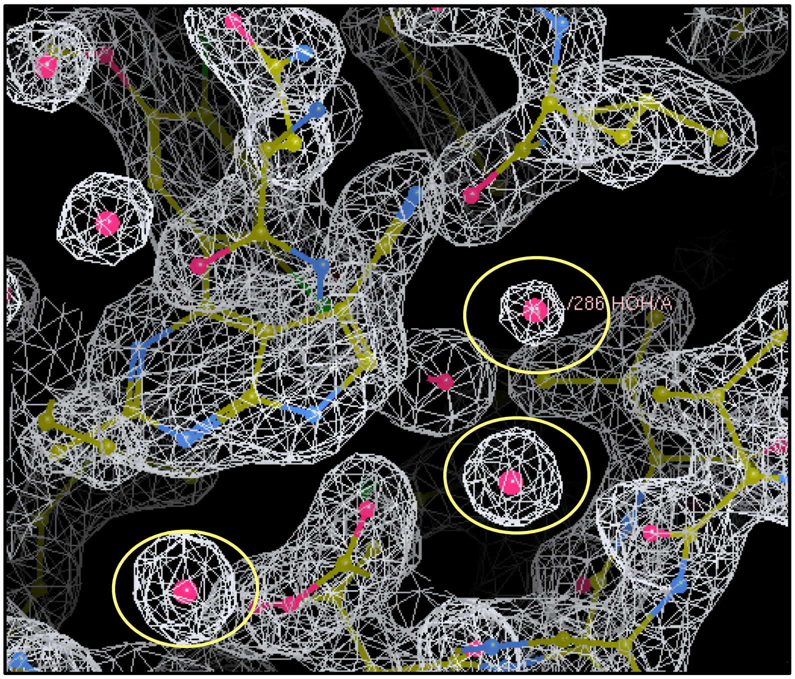


Figure S6: The experimental electron density ($2F_o - F_c$) map (white) of the HSP90 system (PDB: 3RLQ) obtained from https://www.rcsb.org/structure/3RLQ shows a weaker peak for Site 1 compared to Site 2 and 3. The target hydration sites are circled in yellow

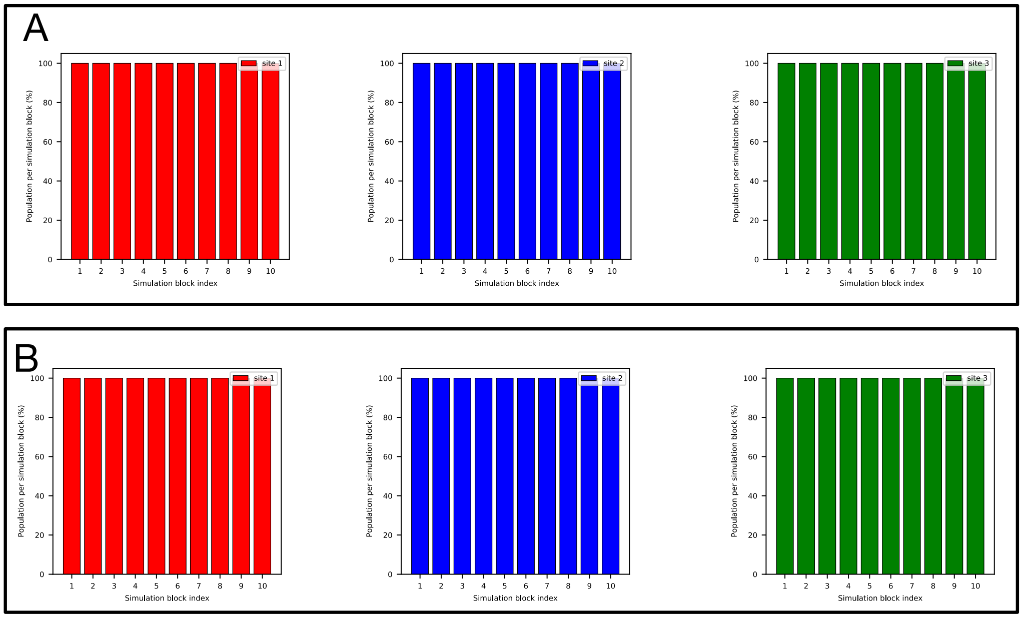


Figure S7: Both BLUES and MD simulations converge to the same occupancy (100%) of all three target sites (Site 1: red, Site 2: blue, Site 3: green) in the HSP90 system (PDB: 2XAB). Bar graphs show the water occupancy of target sites in (A) BLUES simulation and (B) MD simulation. In both simulations, all ordered water molecules were retained prior to simulations.

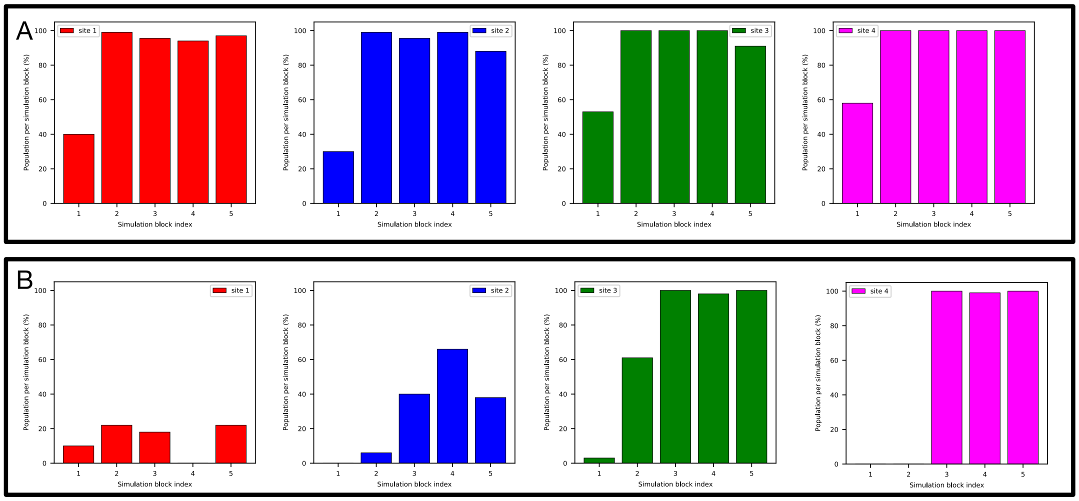


Figure S8: *grand* simulations of the HSP90 system (PDB: 3RLP) do not converge between two replicates in terms of Site 1 and 2 occupancies. Bar graphs show the water occupancy of target sites (Site 1: red, Site 2: blue, Site 3: green, Site 4: magenta) in two replicates of (A-B) *grand* simulations.

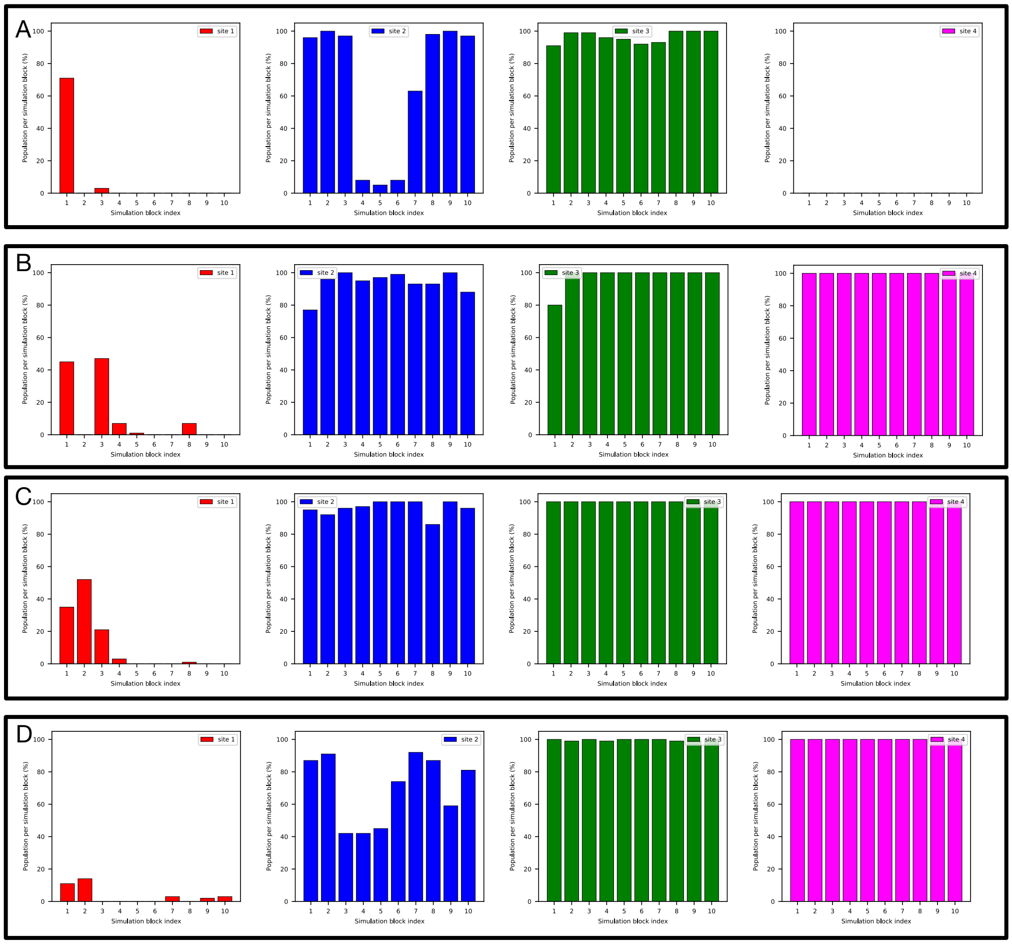


Figure S9: MD simulations suggest a low occupancy of Site 1 in the HSP90 system (PDB: 3RLP). Bar graphs show the water occupancy of target sites (Site 1: red, Site 2: blue, Site 3: green, Site 4: magenta) in MD simulations with ordered water molecules (A-B) removed and (C-D) retained prior to simulations.

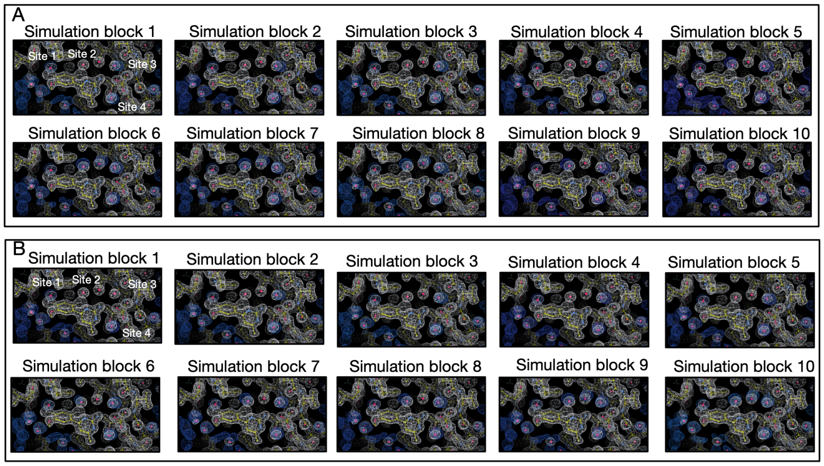


Figure S10: Calculated electron densities (blue) from BLUES simulations agree well with the experimental densities (white) of the HSP90 system (PDB: 3RLP). Target sites are labeled. Panel A and B are results from two replicates.

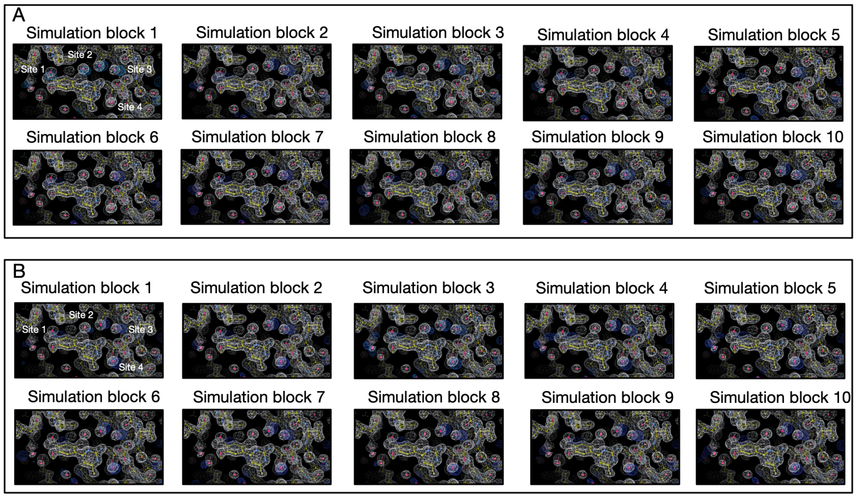


Figure S11: Calculated electron densities (blue) from one replicate of MD simulations agree well with the experimental densities (white) of the HSP90 system (PDB: 3RLP). Target sites are labeled. Panel A and B are results from two replicates.

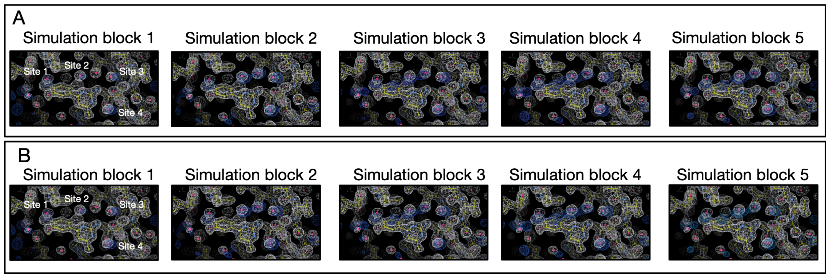


Figure S12: Calculated electron densities (blue) from *grand* simulations agree well with the experimental densities (white) of the HSP90 system (PDB: 3RLP). Target sites are labeled. Panel A and B are results from two replicates.

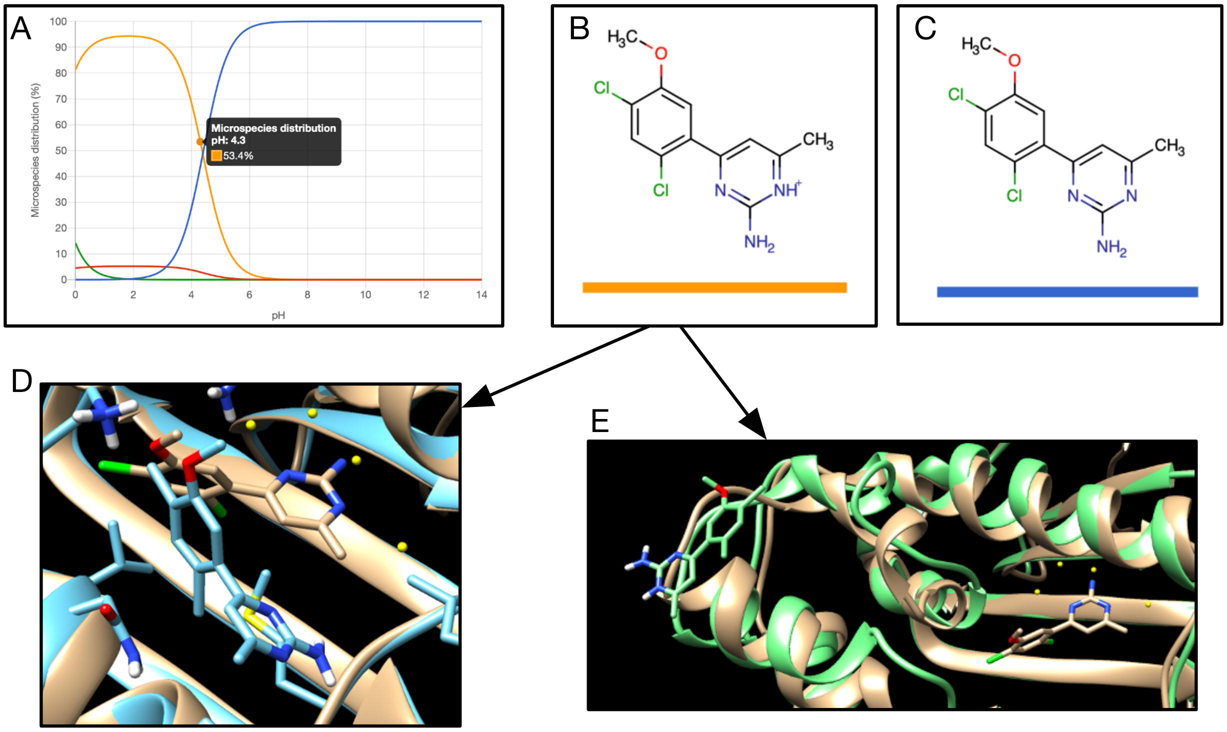


Figure S13: The ligand of the HSP90 system (PDB: 3RLP) has two populated protonation states and we found one of them is not stable in our simulations. (A) Calculated pKa values and the microspecies distribution (in %) of the ligand in the HSP90 system (PDB: 3RLP). (B-C) Populated ligand with different protonation states at pH 4.3. (D-E) Snapshots (blue and green) extracted from simulations using the ligand with protonation state in (B) overlap with the crystallographic pose (tan). The ligand is not stable in the simulation and escapes from the binding site.

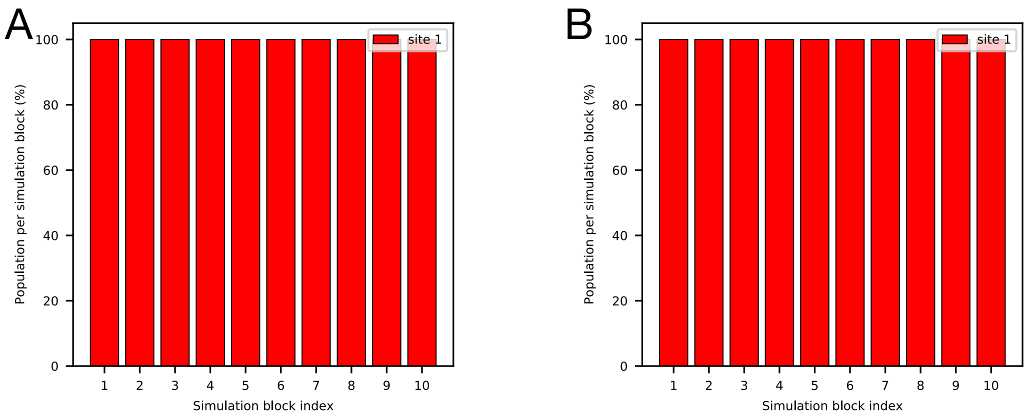


Figure S14: Both BLUES and MD simulations converge to the same occupancy (100%) of the target site (red) in the HSP90 system (PDB: 3RLR). Bar graphs show the water occupancy of target sites of the HSP90 system (PDB: 3RLR) in (A) BLUES simulation and (B) MD simulation. In both simulations, all ordered water molecules were retained in the starting structures.

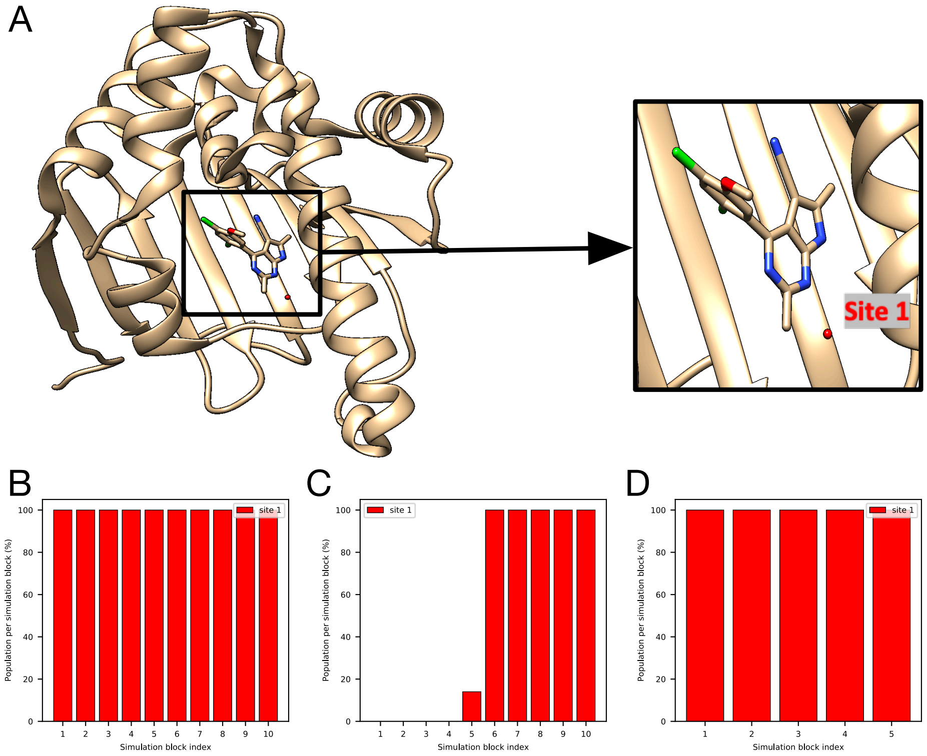


Figure S15: Both BLUES and *grand* simulations converge to the same occupancy (100%) of the target site (red) in (A) the HSP90 system (PDB: 3RLR). Bar graphs show the water occupancy of target sites in a single (B) BLUES simulation (1.4 million force evaluations for each simulation block), (C) *grand* simulation. In both simulations, all ordered water molecules were removed in the starting structures.

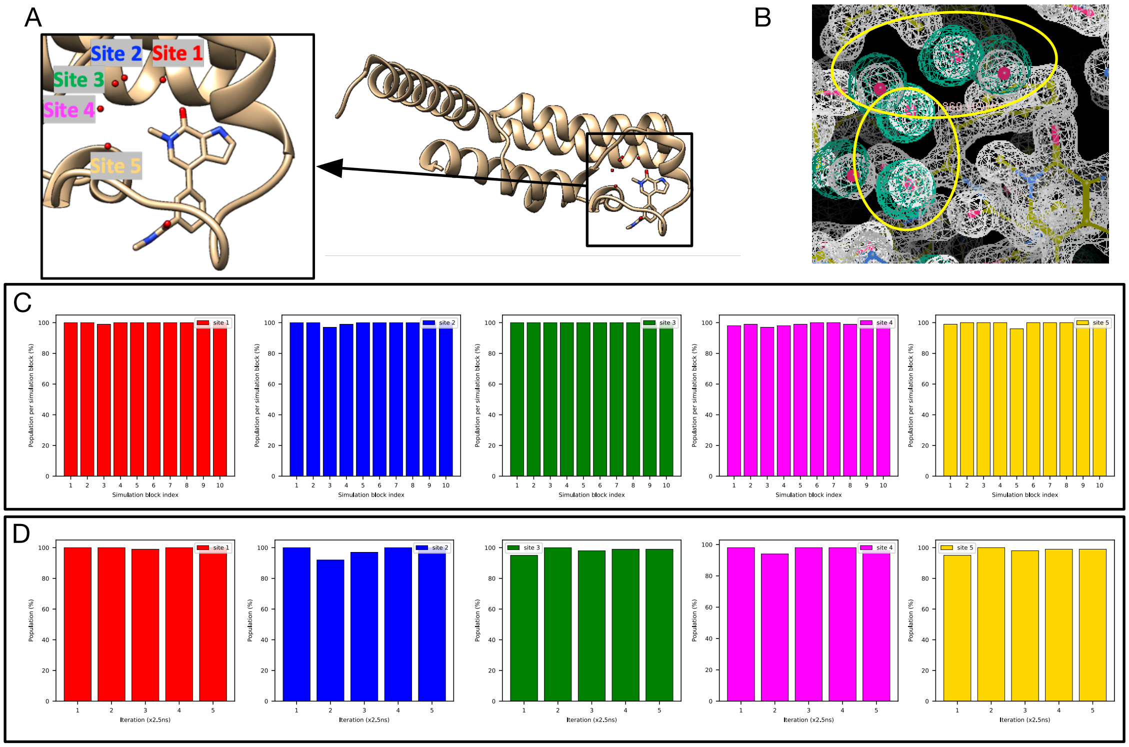


Figure S16: Both BLUES and *grand* simulations converge to the same occupancy (close to 100%) of the target sites (Site 1: red, Site 2: blue, Site 3: green, Site 4: magenta, Site 5: yellow) in (A) The TAF1(2) system (PDB: 5I29). (B) The calculated electron density map (blue) overlaps with the experimental electron density ($2F_o - F_c$) map (white). The target hydration sites are circled. The calculation is based on a BLUES simulation trajectory. Bar graphs show the water occupancy of target sites in a single (C) BLUES simulation, and (D) *grand* simulation.

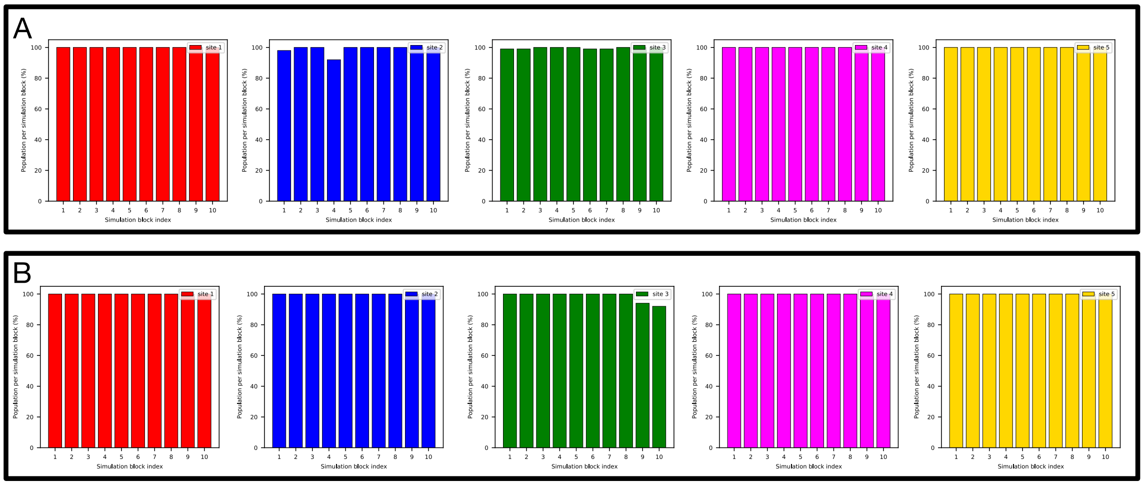


Figure S17: Both replicates of BLUES simulations converge to the same occupancy (close to 100%) of the target sites (Site 1: red, Site 2: blue, Site 3: green, Site 4: magenta, Site 5: yellow) of the TAF1(2) system (PDB: 5I29). Bar graphs show the water occupancy of target sites in BLUES simulations (all ordered water molecules were retained prior to simulations).

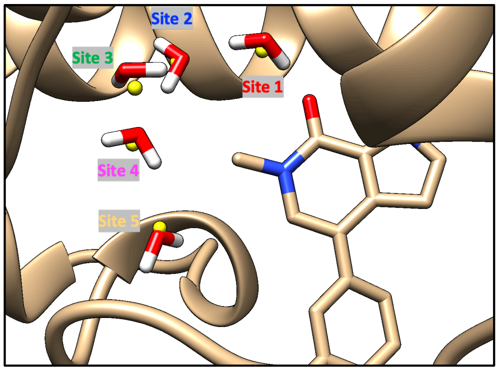


Figure S18: All target sites in the TAF1(2) system (PDB: 5I29) have been rehydrated after equilibration (with position restraints applied). The snapshot is extracted after equilibration simulations. Crystallographic water sites are shown in yellow.

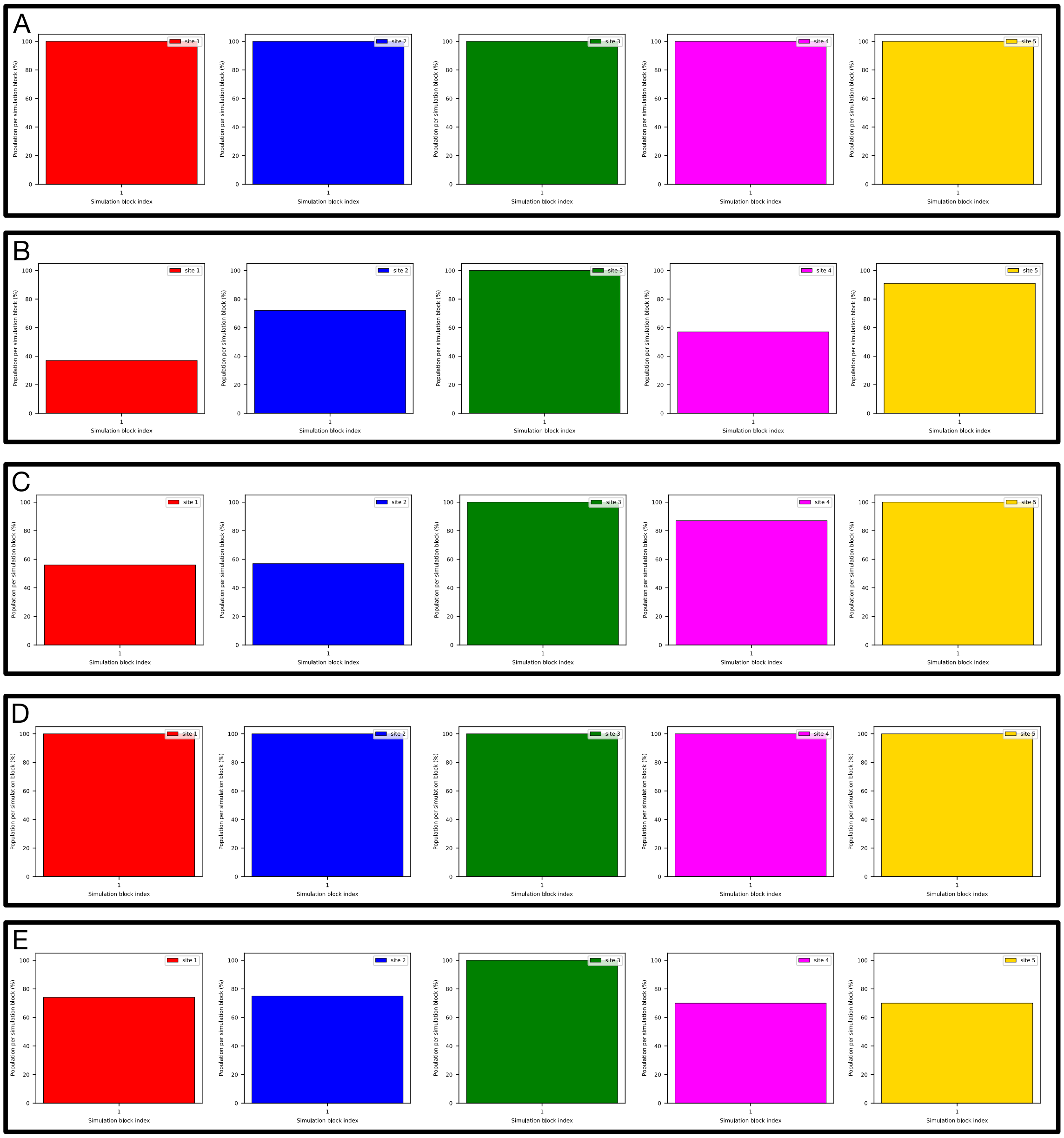


Figure S19: All target sites in the TAF1(2) system (PDB: 5I29) have been rehydrated in most equilibration simulations (with position restraints applied). Bar graphs show the water occupancy of target sites (Site 1: red, Site 2: blue, Site 3: green, Site 4: magenta, Site 5: yellow) in 5 replicates of 10 ns NPT equilibration simulations (all ordered water molecules were removed prior to simulations).

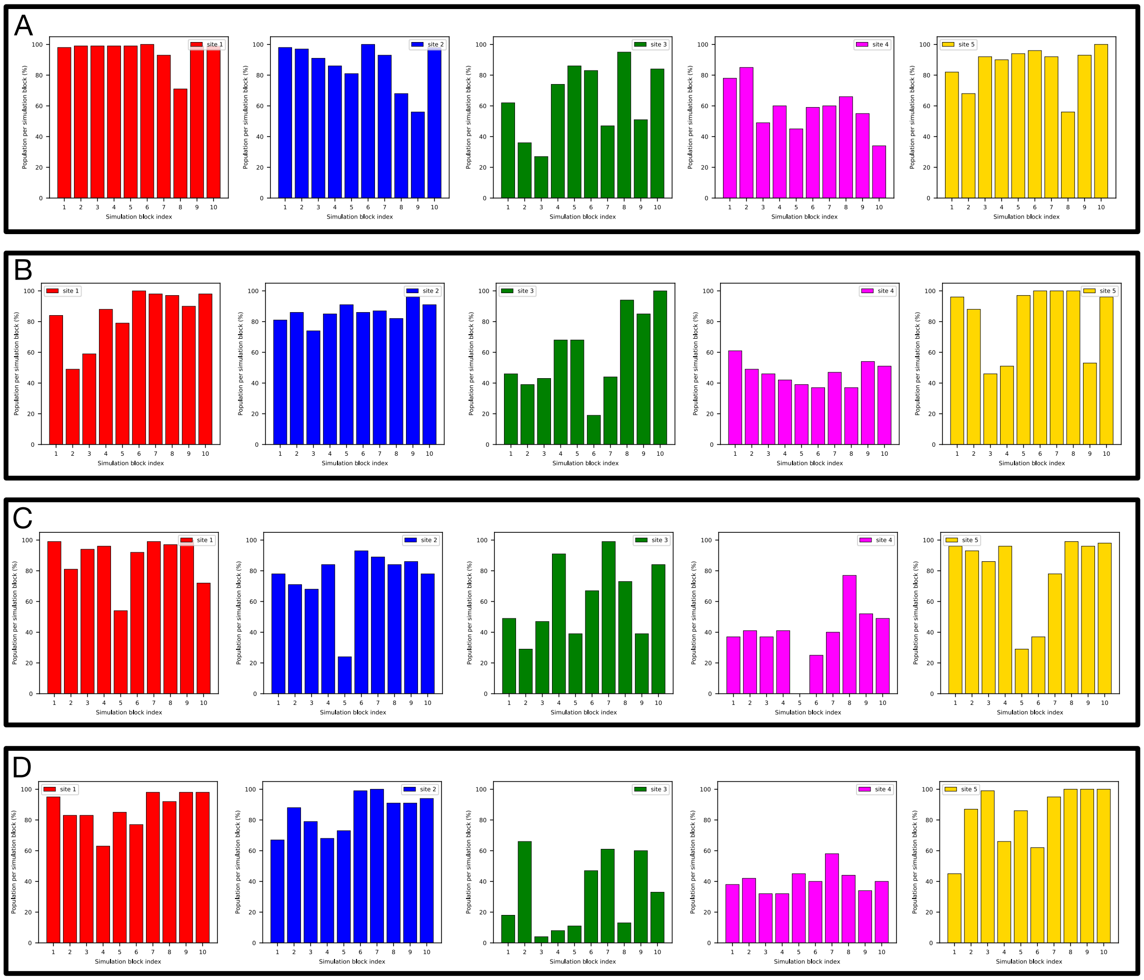


Figure S20: MD simulations of the TAF1(2) system (PDB: 5I29) do not return converged occupancies for target sites especially for Site 3. Bar graphs show the water occupancy of target sites (Site 1: red, Site 2: blue, Site 3: green, Site 4: magenta, Site 5: yellow) in MD simulations with ordered water molecules (A-B) removed and (C-D) retained prior to simulations.

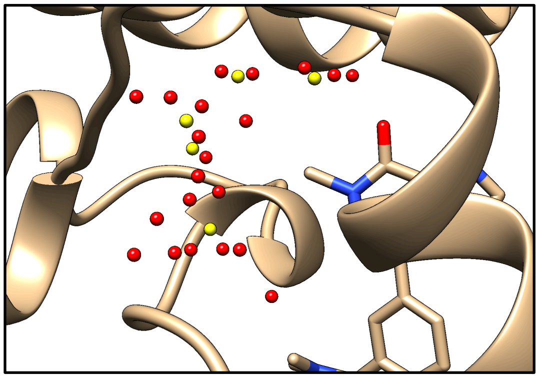

Figure S21: An example of overcrowded water sites from clustering analysis (red) using build-in functions in *grand* package. Crystallographic water molecules are shown in yellow. The crystal structure is shown in tan (TAF1(2), PDB: 5I29).

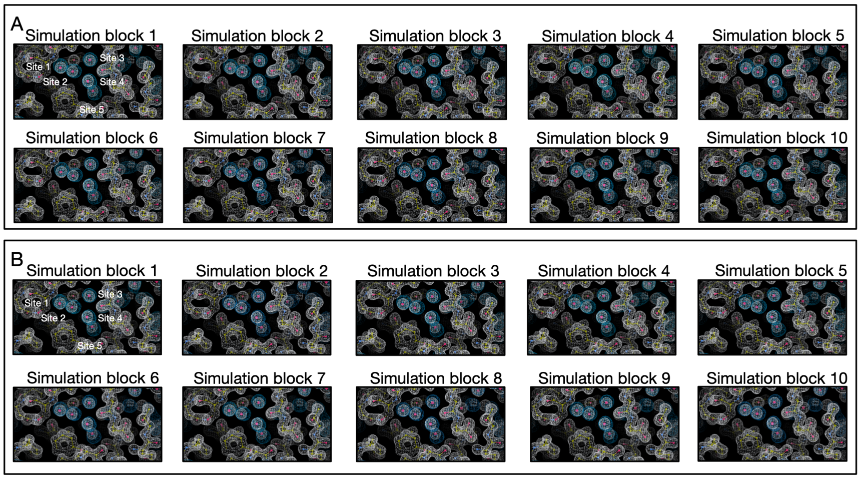


Figure S22: Calculated electron densities (blue) from BLUES simulations agree well with the experimental densities (white) of the TAF1(2) system (PDB: 5I29). Target sites are labeled. Panel A and B are results from two replicates.

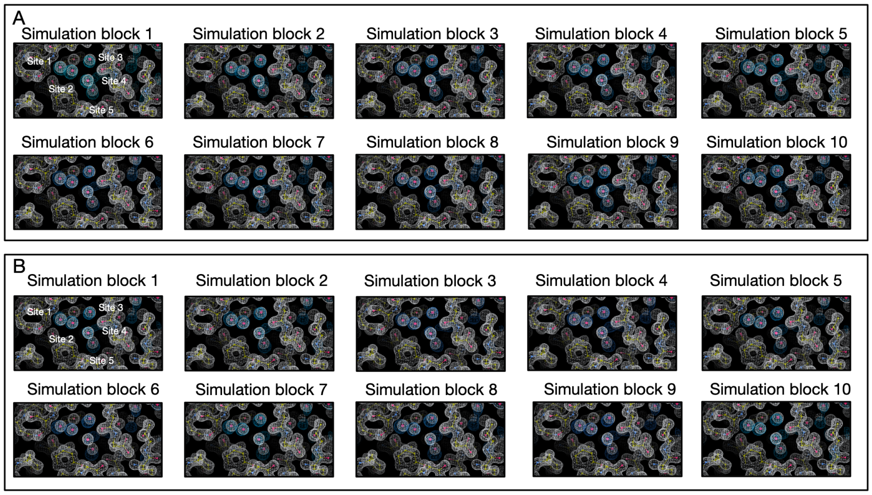


Figure S23: Calculated electron densities (blue) from MD simulations agree well with the experimental densities (white) of the TAF1(2) system (PDB: 5I29). Target sites are labeled. Panel A and B are results from two replicates.

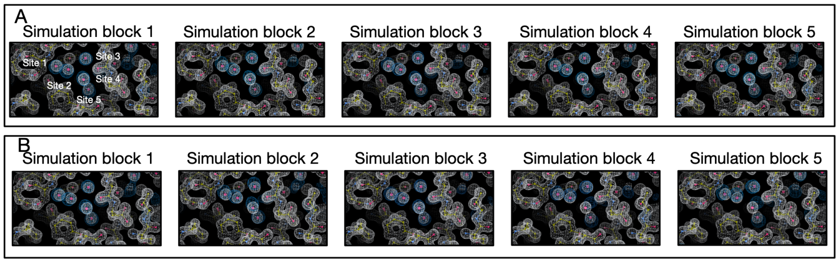


Figure S24: Calculated electron densities (blue) from *grand* simulations agree well with the experimental densities (white) of the TAF1(2) system (PDB: 5I29). Target sites are labeled. Panel A and B are results from two replicates.

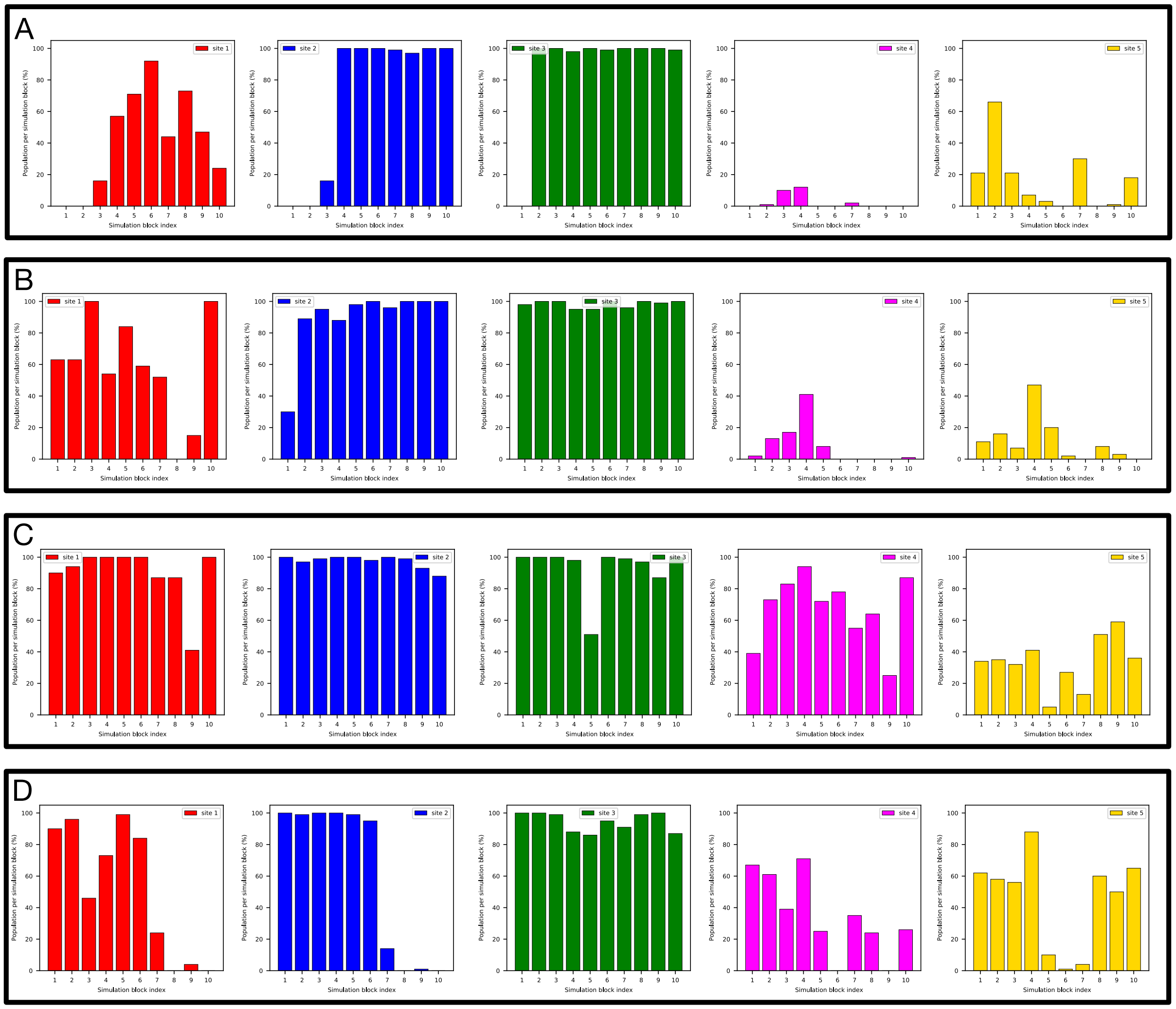


Figure S25: MD simulations do not return converged occupancies between replicates for target sites in the TAF1(2) system (PDB: 5I1Q), especially for Site 4 and 5. Bar graphs show the water occupancy of target sites (Site 1: red, Site 2: blue, Site 3: green, Site 4: magenta, Site 5: yellow) in MD simulations with ordered water molecules (A-B) removed and (C-D) retained prior to simulations.

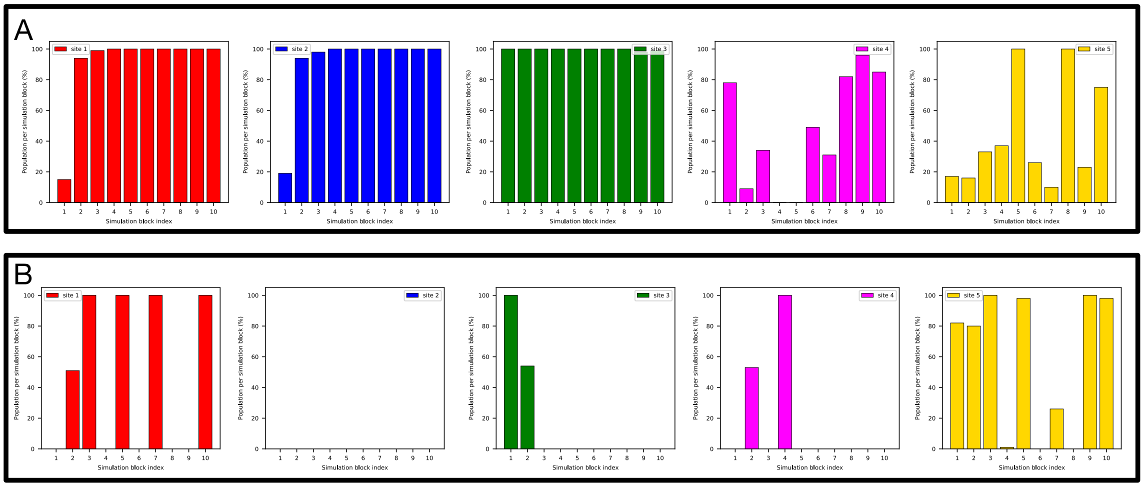


Figure S26: *grand* simulations do not return converged occupancies between replicates of target sites. Bar graphs show the water occupancy of target sites of the TAF1(2) system (PDB: 5I1Q) in *grand* simulations.

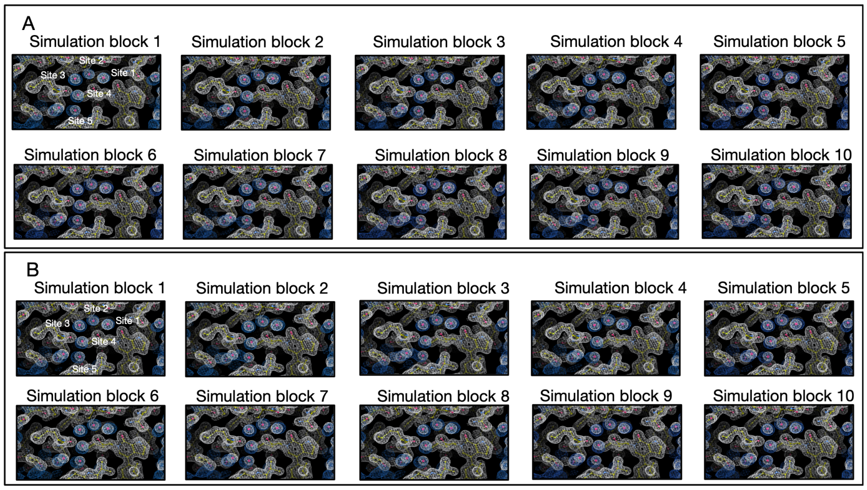


Figure S27: Calculated electron densities (blue) from BLUES simulations agree well with the experimental densities (white) of the TAF1(2) system (PDB: 5I1Q). Target sites are labeled. Panel A and B are results from two replicates.

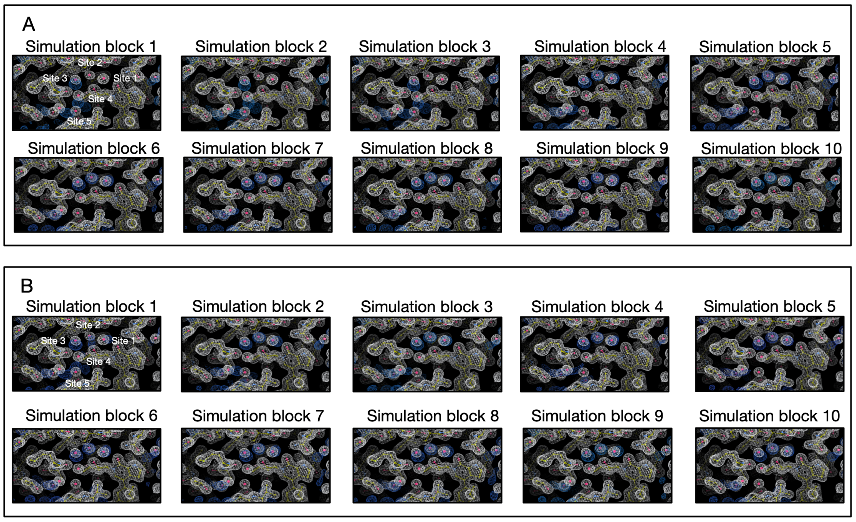


Figure S28: Calculated electron densities (blue) from one replicate of MD simulations agree well with the experimental densities (white) of the TAF1(2) system (PDB: 5I1Q). Target sites are labeled. Panel A and B are results from two replicates.

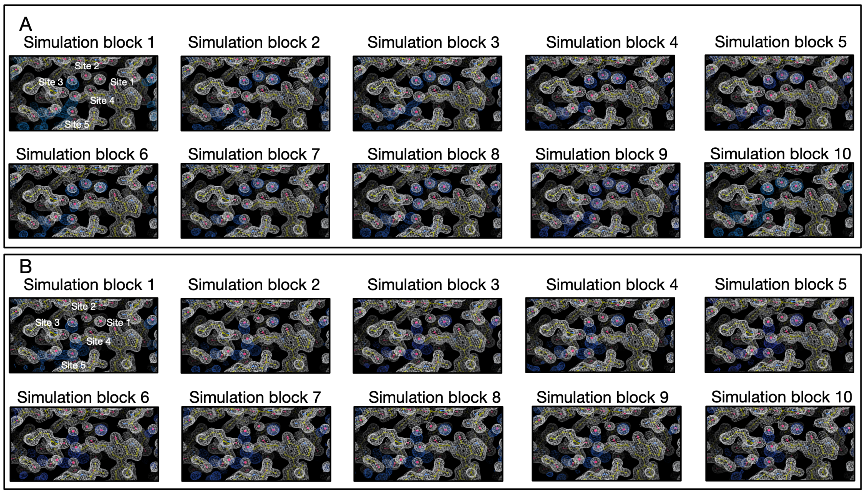


Figure S29: Calculated electron densities (blue) from *grand* simulations agree well with the experimental densities (white) of the TAF1(2) system (PDB: 5I1Q). Target sites are labeled. Panel A and B are results from two replicates.

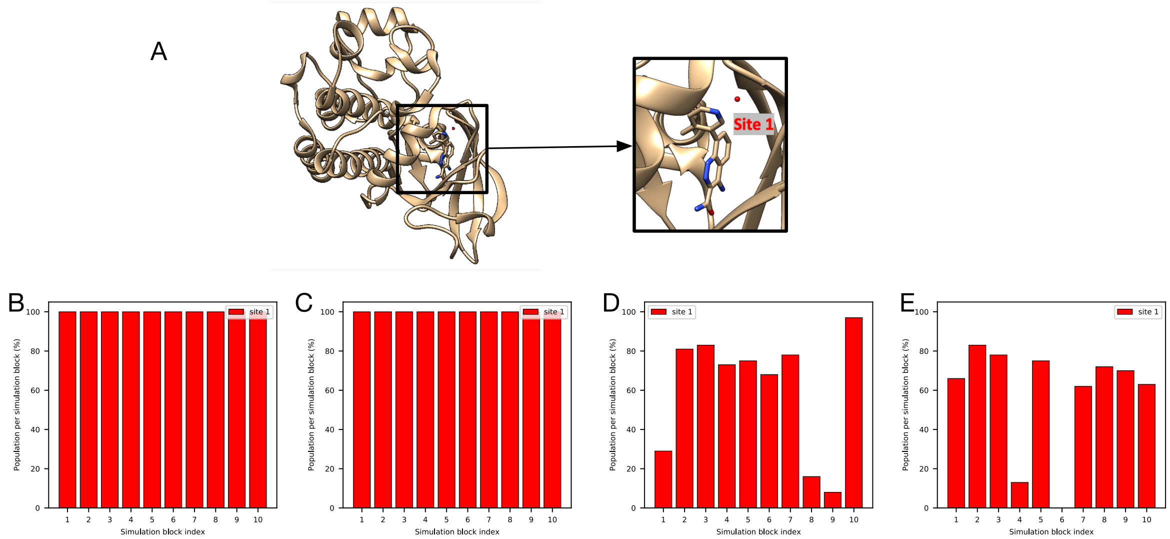


Figure S30: MD and BLUES simulations do not converge to the same occupancy for the target site (red) in (A) the BTK system (PDB: 4ZLZ). Bar graphs show the water occupancy of the target site in (A-B) BLUES simulations and (C-D) MD simulation. In both simulations, all ordered water molecules were retained in the starting structures.

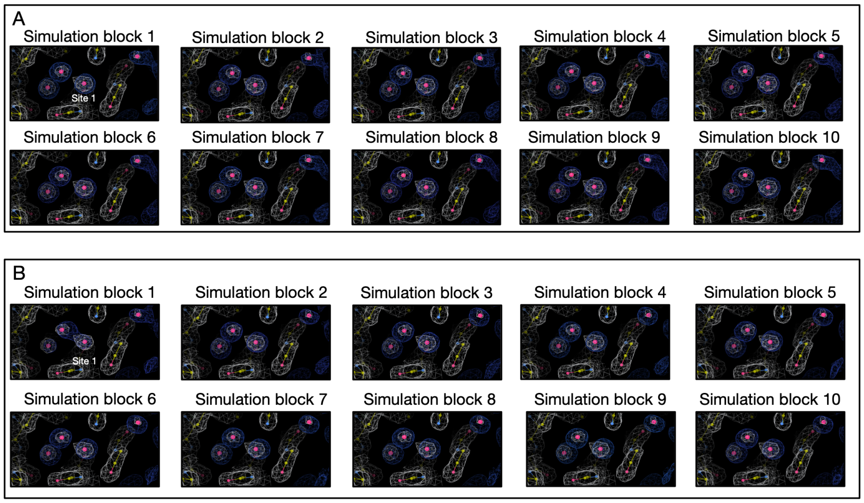


Figure S31: Calculated electron densities (blue) from BLUES simulations agree well with the experimental densities (white) of the BTK system (PDB: 4ZLZ). Target sites are labeled. Panel A and B are results from two replicates.

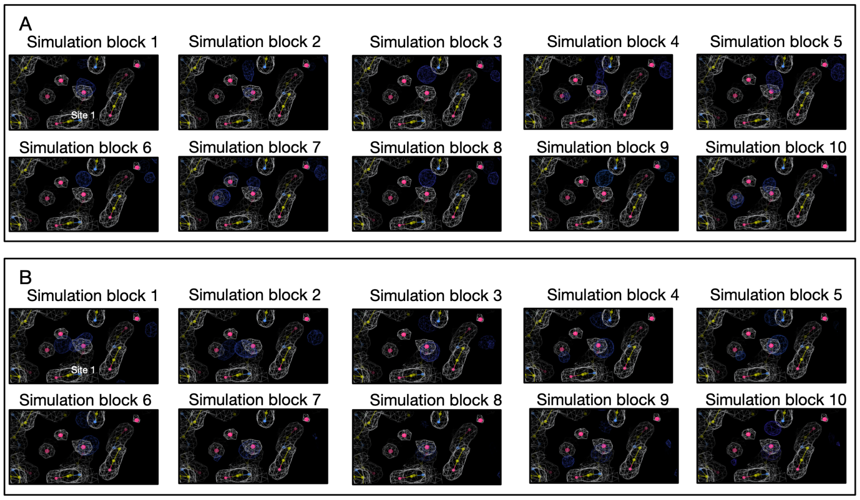


Figure S32: Calculated electron densities (blue) from MD simulations agree well with the experimental densities (white) of the BTK system (PDB: 4ZLZ). Target sites are labeled. Panel A and B are results from two replicates.

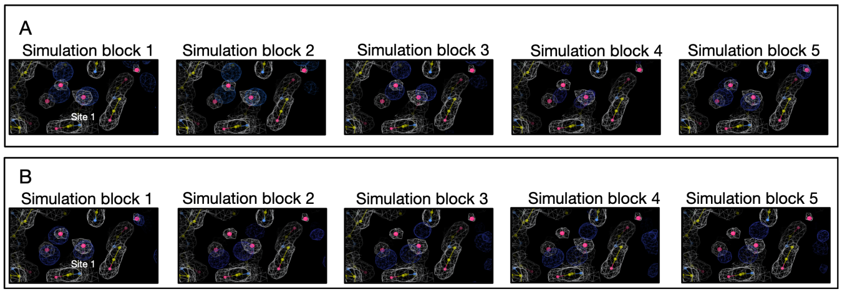


Figure S33: Calculated electron densities (blue) from *grand* simulations agree well with the experimental densities (white) of the BTK system (PDB: 4ZLZ). Target sites are labeled. Panel A and B are results from two replicates.

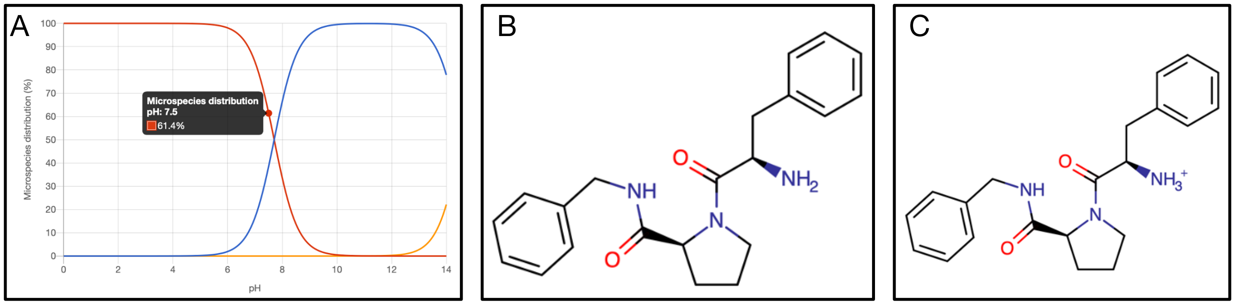


Figure S34: The ligand of the thrombin system (PDB: 2ZFF) has two populated protonation states and we do not observed difference in terms of water site rehydration. (A) Calculated pKa values and the microspecies distribution (in %) of the ligand in the thrombin system (PDB: 2ZFF). (B-C) Populated ligand with different protonation states at pH 7.5.
